# Predicting gene regulatory networks from multi-omics to link genetic risk variants and neuroimmunology to Alzheimer’s disease phenotypes

**DOI:** 10.1101/2021.06.21.449165

**Authors:** Saniya Khullar, Daifeng Wang

**Affiliations:** Department of Biostatistics and Medical Informatics, University of Wisconsin – Madison, Madison, WI 53076, USA; Waisman Center, University of Wisconsin – Madison, Madison, WI 53705, USA; Department of Computer Sciences, University of Wisconsin – Madison, Madison, WI 53706, USA

## Abstract

**Background:** Genome-wide association studies have found many genetic risk variants associated with Alzheimer’s disease (AD). However, how these risk variants affect deeper phenotypes such as disease progression and immune response remains elusive. Also, our understanding of cellular and molecular mechanisms from disease risk variants to various phenotypes is still limited. To address these problems, we performed an integrative multi-omics analysis of genotype, transcriptomics, and epigenomics for revealing gene regulatory mechanisms from disease variants to AD phenotypes.

**Method:** First, given the population gene expression data of a cohort, we construct and cluster its gene co-expression network to identify gene co-expression modules for various AD phenotypes. Next, we predict transcription factors (TFs) regulating co-expressed genes and AD risk SNPs that interrupt TF binding sites on regulatory elements. Finally, we construct a gene regulatory network (GRN) linking SNPs, interrupted TFs, and regulatory elements to target genes and gene modules for each phenotype in the cohort. This network thus provides systematic insights into gene regulatory mechanisms from risk variants to AD phenotypes.

**Results:** Our analysis predicted GRNs in three major AD-relevant regions: Hippocampus, Dorsolateral Prefrontal Cortex (DLPFC), Lateral Temporal Lobe (LTL). Comparative analyses revealed cross-region-conserved and region-specific GRNs, in which many immunological genes are present. For instance, SNPs rs13404184 and rs61068452 disrupt SPI1 binding and regulation of AD gene INPP5D in the Hippocampus and LTL. However, SNP rs117863556 interrupts bindings of REST to regulate GAB2 in DLPFC only. Driven by emerging neuroinflammation in AD, we used Covid-19 as a proxy to identify possible regulatory mechanisms for neuroimmunology in AD. To this end, we looked at the GRN subnetworks relating to genes from shared AD-Covid pathways. From those subnetworks, our machine learning analysis prioritized the AD-Covid genes for predicting Covid-19 severity. Decision Curve Analysis also validated our AD-Covid genes outperform known Covid-19 genes for classifying severe Covid-19 patients. This suggests AD-Covid genes along with linked SNPs can be potential novel biomarkers for neuroimmunology in AD. Finally, our results are open-source available as a comprehensive functional genomic map for AD, providing a deeper mechanistic understanding of the interplay among multi-omics, brain regions, gene functions like neuroimmunology, and phenotypes.

## Introduction

Alzheimer’s Disease (AD), a neurodegenerative disease, affects over 50 million elders worldwide^1^. Late-onset AD (LOAD) comprises >97% of all cases, usually occurring after age 65^2^. AD patients experience phenotypic changes such as memory loss, cognitive decline, and weak executive function (e.g., poor Mini-Mental State Examination scores)^1^. Also, many underlying molecular changes happen, like amyloid-beta plaques, neurofibrillary tangles (NFTs), and chronic neuroinflammation (by dysregulated innate immunity). However, molecular mechanisms causing AD progression and phenotypes remain elusive.

Gene expression and regulation is a key mechanism leading to human diseases. Studies have found differentially expressed genes (DEGs) in AD in different brain regions, e.g., Hippocampus CA1, Lateral Temporal Lobe (LTL), Dorsolateral Prefrontal Cortex (DLPFC), implying important roles of those regions to AD. Also, gene co-expression networks have been widely used to identify co-expressed gene modules and link gene expression patterns to AD phenotypes^3^. Genes in a module show similar expression dynamics across AD phenotypes, implying they share certain molecular mechanisms dysregulated in AD^4^. Nevertheless, understanding gene regulatory mechanisms controlling those DEGs and co-expressed genes and modules for various AD phenotypes is unclear.

Recent Genome-Wide Association Studies (GWAS) identified many genetic variants associated with AD^5^. Linking those AD variants to genes and regulatory mechanisms provides a deeper understanding of molecular causes in AD. AD SNPs have further linked several AD genes, (e.g. BIN1, MS4A4A, SPI1, TP53INP1^6^) plus 11 more by Proteome-Wide Association Studies^7^. TREM2 and CD33 mutations are associated with microglia (resident macrophage immune cells of the brain) activation, increased cytokines, and higher AD risk^8^. However, most AD SNPs locate on non-coding DNA, and it is still challenging to link them to potential AD genes.

Biologically, gene expression is controlled by different regulatory factors such as regulatory variants, transcription factors (TFs), and regulatory elements (e.g., enhancers, promoters). Those factors work together as a gene regulatory network (GRN) to control the expression and functions of genes. For instance, a SNP can disrupt binding sites of TFs (TFBSs) on enhancers to cause abnormal gene expression, potentially leading to diseases, a.k.a, potential disease genes. Using such GRNs, recent studies have predicted many disease genes and functions driven by disease risk variants, such as 321 high-confident Schizophrenia genes^9^. However, how GRNs link AD variants to genes and functions for phenotypes is unclear, especially across regions.

To address these issues, we performed an integrative analysis of multi-omics to reveal genes, functions, and GRNs from AD variants to AD phenotypes for three brain regions as above (**Fig. 1, Methods**). Given a region, we first build a gene co-expression network using population gene expression data from an AD cohort and identify co-expressed genes and modules associated with AD phenotypes. We then integrate chromatin interaction data (e.g., Hi-C) and TF-gene expression relationships to predict TFs regulating co-expressed genes if they bind to the regulatory elements that control them. Finally, we find AD SNPs interrupting those TFBSs, and output a brain-region GRN linking them together. Moreover, since altered immunity is a major factor contributing to LOAD (e.g., neuroinflammation could start decades before), we used those GRNs to link potential biomarker genes and variants for neuroimmunology in AD.

**Fig 1.**
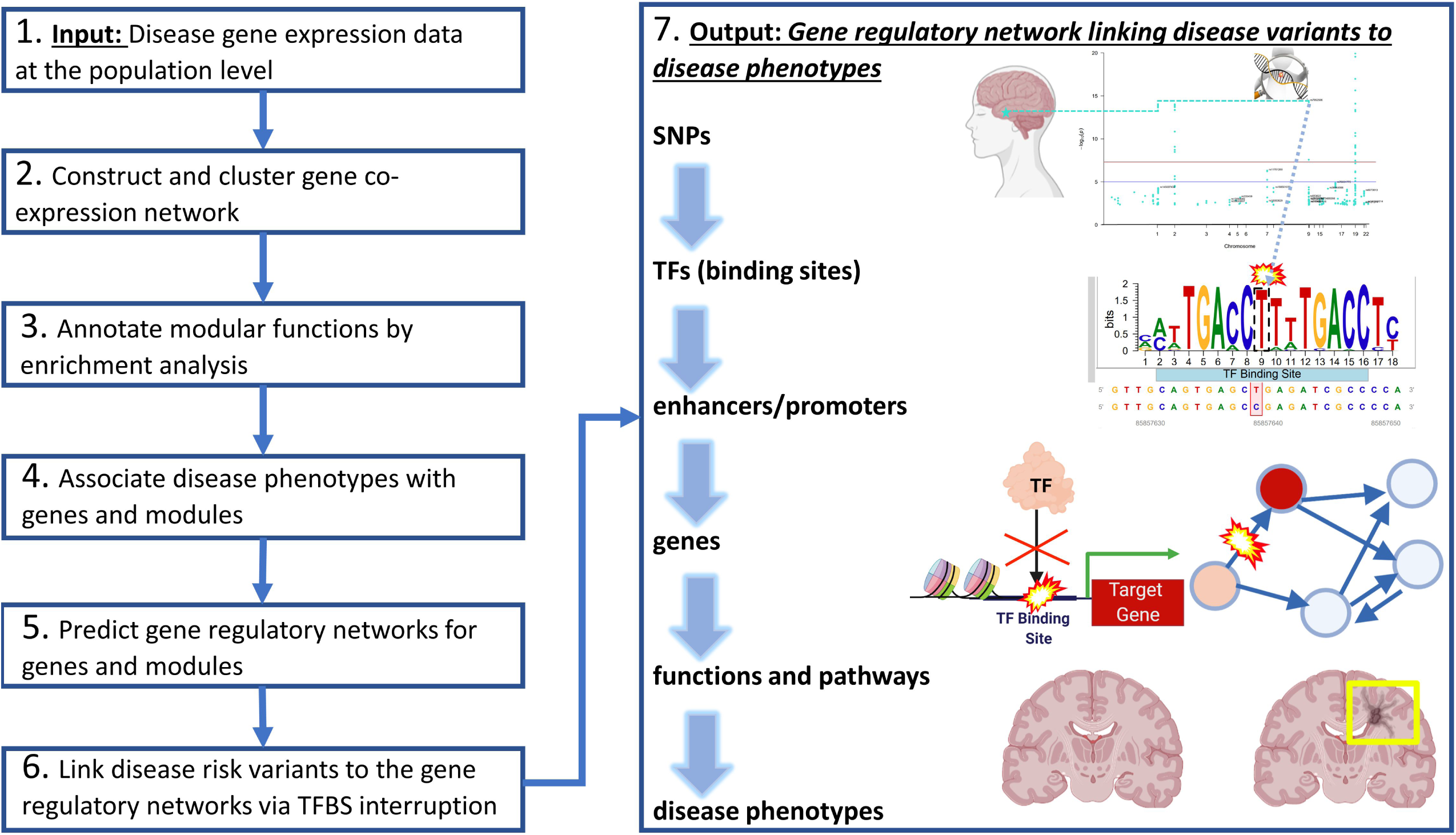
Integrative analyses to predict gene regulatory networks from disease risk variants to phenotypes. Primarily, this analysis consists of seven major steps as a pipeline. First, it inputs the population gene expression data with phenotypic information (Step 1) and constructs and clusters gene co-expression networks into gene modules (Step 2). Second, it performs enrichment analysis for modules (Step 3) and links genes and modules to various phenotypes from the population (Step 4). Third, it predicts the transcription factors (TFs) and regulatory elements (e.g., enhancers) that regulate genes and co-regulate modular genes as a gene regulatory network (Step 5). Also, it further finds disease risk variants (e.g., GWAS SNPs) that interrupt the binding sites of TFs from the network (Step 6). Finally, we output a full gene regulatory network linking disease variants to interrupted TFs and enhancers to regulated genes and modules to enriched functions and pathways to disease phenotypes (Step 7). The network thus provides a deeper understanding of gene regulatory mechanisms in diseases. As a demo, in this paper, we applied to AD population datasets from different brain regions. We predicted brain-specific gene regulatory networks for various AD phenotypes such as progression stages.

Particularly, we looked at the genes from the neuroimmunology pathways shared between AD and SARS-CoV-2 virus (Covid-19), two diseases with a recently reported strong correlation. AD brains have an increased level of circulating cytokines, associated with the cytokine storm (overactivated inflammatory immune system response) impacting Covid-19 morbidity and mortality^10^. Covid-19 has widely affected elders with neurodegenerative diseases and can mediate neuroinflammation^11^, thus providing potential novel insights on a dysregulated immune system in AD and progression. Covid-19 survivors can also have an increased risk of neurological and psychiatric problems, known as Neuro-COVID^12^. β-coronaviruses (like Covid-19) may attack the Central Nervous System, elevating AD dementia processes in the brain^13^. On the other hand, AD patients have a threefold higher risk of infection and doubly higher risk of death from Covid-19^14^. AD risk genes, like APOE4, increase severe Covid-19 susceptibility two-fold^15^. Therefore, looking at gene regulatory mechanisms for shared AD-Covid pathways potentially enables a better mechanistic understanding of neuroimmunology in AD.

The NFKB pathway, shared by AD and Covid-19, regulates normal brain homeostasis (maintaining synaptic plasticity, neurite outgrowth, learning, and memory; moderating neuronal survival and apoptosis; processing neuronal information)^10^, innate immunity and cellular processes including inflammation and cell growth^16^. Since Covid-19 serves as a strong marker for an exaggerated and overreactive immune system, elucidating pathways disrupted in both Covid-19 and AD may provide more insights on misguided immunity in AD onset and progression. A prominent hypothesis believes AD is primarily caused by impaired NFKB pathway signaling, where NFKB TFs are activated, resulting in neuroinflammation, oxidative stress complications, microglia activation, increased cytokine levels, and neuronal death observed in AD brain regions^10^. NFKB TFs are also involved in a positive feedback loop with activating cytokines during Covid-19, contributing to exaggerated inflammatory responses in severe Covid-19^17^. Along with the NFKB Pathway (which induces expression of proinflammatory cytokines and is a key driver in both diseases), there are several shared pathways in Covid-19 and AD. However, underlying gene functions linking immunological functions from Covid-19 to AD are unknown. Finally, we found the subnetworks of GRNs relating to genes from AD-Covid shared pathways. We then trained a machine learning model to predict severe Covid-19 from those network genes using RNA-seq data of a recent Covid-19 cohort. Our network genes outperform known Covid-19 genes for prediction. Using our model, we prioritized a set of highly predictive genes as AD-COVID genes, which can be potential biomarkers for immune system dysregulation and inflammation in AD (since Covid-19 severity is correlated with excessive proinflammatory immune response^18^).

## Materials and Methods

### The pipeline of our integrative analysis for predicting gene regulatory mechanisms from AD risk variants to phenotypes

Our analysis can be summarized as a pipeline to predict gene regulatory networks (GRNs) from disease risk variants to phenotypes (**Fig. 1**). The network for specific phenotypes links disease risk variants (e.g., GWAS SNPs), non-coding regulatory elements, transcription factors (TFs) to genes and genome functions, providing comprehensive mechanistic insights on gene regulation in disease phenotypes. Specifically, the pipeline includes the following steps. Here, our analysis is open-source available at https://github.com/daifengwanglab/ADSNPheno.

- Step 1: Input gene expression data at the population level. The input data includes gene expression data of individuals and clinical information on AD phenotypes such as Braak staging and progression.
- Step 2: Construct and cluster gene co-expression network. The pipeline constructs a gene co-expression network linking all possible gene pairs from the input data. The network edge weights are the Pearson correlations of the gene expression profiles across input samples. The gene co-expression network is further clustered into gene co-expression network modules. The genes in the same co-expression module are likely involved in similar functions and co-regulated by specific regulatory mechanisms.
- Step 3: Annotate modular functions by enrichment analyses. To annotate the functions of gene co-expression modules, we calculate enriched pathways and functions, including KEGG pathways, REACTOME pathways, and Gene Ontology (GO) terms of the genes in each gene co-expression module.
- Step 4: Associate AD phenotypes with genes and modules. We associate genes and modules with the phenotypes of input samples, revealing potential driver genes and modules for the phenotypes.
- Step 5: Predict GRNs for genes and modules. We apply multiple computational methods to predict the gene regulatory networks that link TFs, non-coding regulatory elements to genes and modules, providing regulatory mechanistic insights on AD genes and modules.
- Step 6: Link disease risk variants to the gene regulatory network. Our analysis further finds disease risk variants that interrupt the binding sites of TFs (TFBSs) in the gene regulatory networks for identifying functional variants to genes and modules to AD phenotypes.
- Step 7: Output a gene regulatory network linking disease variants to AD phenotypes. Ultimately, this network is the output that links AD genetic risk variants, non-coding regulatory elements, TFs to genes, and genome functions (via modules) for various phenotypes in the input data.

### Population gene expression data and data processing in Alzheimer’s disease

We applied this pipelined analysis to post-mortem human AD population gene expression datasets for three major AD brain regions: Hippocampal CA1, Lateral Temporal Lobe (LTL), and Dorsolateral Prefrontal Cortex (DLPFC). Also, we processed the datasets as follows.

Hippocampal CA1: The microarray gene expression dataset (GSE1297)^19^ was used. The dataset had total RNA expression values for 22,283 HG-U133 Affymetrix Human Genome U133 Plus 2.0 Microarray Identifier probes for 31 individual samples. These individual samples include 9 control (no AD), 7 initial stage, 8 moderate stage, and 7 severe stage samples. We used GEOquery^20^, hgu133a.db^21^, hgu133acdf^22^, and Affy^23^ R packages to download raw data and perform Robust Multichip Average (RMA) normalization^24^ to account for background and technical variations among the samples. We mapped microarray probes to genes, averaging values that mapped to the same gene Entrez ID, removing unmapped probes. We transformed the resulting data by log2(x + 1) transformation and standardized that by scale() function in R. The final Hippocampal CA1 gene expression dataset has 13,073 unique genes for 31 samples.

Lateral Temporal Lobe (LTL): The normalized bulk RNA-Seq dataset (GSE159699)^25^ was used. This dataset had total RNA expression values for 27,130 different genes for 30 individual samples. This group of individual samples includes 8 young (below age 60),10 old, and 12 old samples with advanced AD. After our data pre-processing steps, we had 25,292 genes, and we applied a log2(x+1) transformation to this data. The finalized gene expression dataset in Lateral Temporal Lobe has 25,292 unique genes for these 30 samples.

Dorsolateral Prefrontal Cortex (DLPFC): FPKM data from the ROSMAP Study, available on synapse.org (ID: syn3219045), was used^26^. Removing lowly expressed protein-coding genes (those with 0 variance and relative weights below 0.1), using WGCNA’s goodSamplesGenes() function, shrunk the list of DLPFC genes down to 26,014 genes. In this dataset, there are 638 out of 640 individual RNA-Seq samples with mapped phenotype information. We also applied a log2(x+1) transformation to this gene expression data and then standardized it with R’s scale() function. The finalized DLPFC gene expression dataset has 26,014 genes for 638 samples.

Finally, there are 12,183 shared genes across these 3 brain regions (Venn Diagram in **Fig. S1**).

### Regulatory elements and Chromatin interactions in the human brain regions

Epigenomic data has identified a variety of regulatory elements such as enhancers and promoters. Chromatin interaction data (e.g., Hi-C) have further revealed interacting enhancers and gene promoters. Thus, we integrated recent published epigenomic and chromatin interaction data for three brain regions to link enhancers to genes (via promoters). For Hippocampal Ca1, we obtained its enhancers and promoters from Brain Open Chromatin Atlas^27^ and promoter-based interactions from GSE86189^28^. To identify promoters in LTL and DLPFC, we used R package, TxDb.Hsapiens.UCSC.hg19.knownGene^29^, to retrieve promoter start and stop positions of genes, using a short ultra-conserved promoter length of 5,000 base pairs upstream of the protein-coding start site^30^. Besides, we used the H3K27ac data from GSE130746^25^ to find LTL enhancers. This dataset contains information on the target gene, distance that the H3K27ac mark is from the target gene’s Transcription Start Site (TSS), and enhancer start and end positions. The enhancers in LTL that we used were at least 1,000 bases away from the TSS. Moreover, for DLPFC, we used the enhancers and interacting enhancer-promoter pairs in DLPFC from PsychENCODE^31^.

### Gene co-expression network analysis

We applied WGCNA^32^ to population gene expression data to construct and cluster gene co-expression networks into gene co-expression modules (minimum module size = 30 genes). Also, we further used a K-Means clustering step based on code^33^ and methodology that has been previously applied to conventional WGCNA and proven to improve module assignments and functional enrichments, in applications such as for brain-specific cell-type marker enrichments^34^. This additional K-means step utilizes the modular eigengenes (MEs) from WGCNA modules as initial centroids, initial gene assignments, and computable distance between gene and MEs for K-Means to re-assign genes to optimal modules across the iterations. In total, we obtained 30 gene co-expression modules for Hippocampus (13,073 genes), 56 modules for the LTL (25,292 genes), 35 modules for DLPFC (26,014 genes).

### Enrichment analyses of gene co-expression modules

Co-expressed genes in the same module are highly likely involved in similar functions and pathways. The enrichment analysis has thus been widely used to identify such functions and pathways in a gene module. Enrichment p-values were adjusted using the Benjamini-Hochberg (B-H) correction. Given a group of genes (e.g., from a module) for each brain region, we used multiple tools and hundreds of data sources for enrichment analyses (**Table S1**). We used the highest enrichment -log10(adjusted p-value) scores from any source for each gene module and respective enrichment in a brain region. Then, for each enrichment for a phenotype in a region, we averaged the non-zero enrichment values for the gene modules that are statistically significantly positively correlated for that phenotype.

### Association of genes and modules with AD phenotypes

We further associated genes and modules with these key AD developmental phenotypes: AD Stages and Progression (Moderate Stage, Severe Stage, and AD Progression), Healthy/Resilient (Control Stage or other resilient individuals with better cognitive abilities despite AD pathology), APOE genotype (APOE E4/E4 is a huge AD risk factor^35^), Braak Staging, neuritic plaque accumulation (measured by CERAD Score), and cognitive impairment level (based on the MMSE Score). We associated gene co-expression modules with all possible AD phenotypes from the input data, by computing the correlations of each modular eigengene (ME) to each phenotype. The eigengenes of modules by WGCNA are the first principal components of modular gene expression. A ME is a vector representing expression levels of input samples and is the likeliest gene expression pattern of module genes. Second, based on the MEs, we used moduleTraitCor() and moduleTraitPvalue() in WGCNA to correlate our MEs with our phenotypes and find the most significantly associated phenotypes to the modules. Statistically significant module-phenotype associations for analysis have a p-value less than 0.05 and a positive correlation. Also, we used gene co-expression networks to examine the relationship between genes and AD phenotypes and identify potential driver (hub) genes for the modules (based on degree of connectivity for each gene in each module).

### Prediction of gene regulatory networks

Gene Regulatory Networks (GRNs), a key molecular mechanism, fundamentally control gene expression. Also, co-expressed genes are likely co-regulated by similar GRNs. Thus, our analysis integrates multiple methods to predict GRNs from gene expression data and co-expression modules. This study predicted GRNs in 3 brain regions that link transcription factors (TFs), regulatory elements, and target genes/modules. First, we identified regulatory elements (e.g. enhancers and promoters) that potentially interact using recent chromatin interaction data (Hi-C) and the scGRNom pipeline^36^. Second, we inferred the TF binding sites (TFBSs) based on consensus binding site sequences on interacting enhancers and promoters by TFBSTools^37^ and motifmatchr^38^. This step generates a reference network linking TFs to regulatory elements (by TFBSs) to genes (by interactions). Third, using gene expression data for a given brain region, we predicted all possible TF-target gene (TG) pairs (or TF-modules) that have strong expression relationships by 3 widely used tools and databases: RTN^39^, TReNA Ensemble Solver^40^, Genie3^41^, (and TF-gene-module pairs by RTN), as below. Finally, this step maps these TF-TG pairs to the reference network. It outputs a full GRN for the region that links TFs, non-coding regulatory elements to target genes and modules.

We combined a recent list of TFs^42^ with JASPAR’s list^43^ to generate a final list of candidate TFs for inferring TF-TG pairs with strong expression relationships. We used this final TF list to find candidate TFs for each brain region (based on genes in the respective gene expression data). Also, TReNA Ensemble Solver with the default parameters (geneCutoff of 0.1 and ensemble of these solvers: LassoSolver, RidgeSolver, RandomForestSolver, LassoPVSolver, PearsonSolver, and SpearmanSolver) was used to construct the transcriptional regulatory network that links TFs to target genes (TGs). Besides, we used GENIE3 to predict additional GRNs via Random Forest regression, predicting each gene’s expression pattern from the expression patterns of all TFs (TF-TG pairs with weights greater than 0.0025 were retained). In addition, we used RTN to predict TFs to TGs by calculating the Mutual Information between the TFs and all genes; permutation analysis with 1,000 permutations, bootstrapping, and the ARACNe algorithm^44^ were used to select most meaningful network edges. Finally, the TF-TG pairs found in at least 2 of the above 3 sources were combined to map to the reference network. For the DLPFC, we instead used the published PsychENCODE GRN (Elastic Net regression weight of 0.1 as a cutoff) filtered for genes found in the DLPFC gene expression data^9^.

In addition to predicting TFs for individual genes, we inferred TFs significantly co-regulating genes in a module in the Hippocampus and LTL. In particular, we performed Master Regulatory Analysis (MRA) on the RTN-inferred network by the RTN package^39^. For each gene module, MRA performed enrichment analysis using the inferred GRN, the phenotype (Module Membership correlation of all genes to that module), and hits (genes assigned to that module). It applied the hypergeometric test for overlaps between TFs and the genes (using gene expression data) and found the statistically significant TFs for each module.

### Linking GWAS SNPs for AD to gene regulatory elements

GWAS studies have identified a wide variety of genetic risk variants associated with diseases. However, most disease risk variants lie on non-coding regions, hindering finding disease genes and understanding downstream disease functions. To this end, we used GRNs as described above to link AD SNPs to regulatory elements in the networks via interrupted TFBSs. We looked at 97,058 GWAS SNPs significantly associated with AD (i.e., AD risk SNPs with p<0.005)^5^. We overlapped those AD risk SNPs with the regulatory elements such as enhancers and promoters in the GRNs from Step 5. Then, we identified the variants that interrupt the TFBSs on the regulatory elements by motifbreakR^45^ (using ENCODE-motif, FactorBook, HOCOMOCO, and HOMER data sources, default methodology, threshold of 0.001), and further linked them to the genes from the regulatory elements with interrupted TFBSs. An extension of 10 kilobase pairs was added to start and end positions of enhancers and an extension of 2 kilobase pairs was added to start and end positions of promoters. We found that 83,842 SNPs out of 97,058 AD GWAS SNPs interrupt the binding sites of 787 TFs.

### Identification of AD-Covid genes and regulatory networks

To investigate potential mechanistic interplays between AD and Covid-19, we compared KEGG pathways^46^ for AD (hsa05010) and Covid-19 (hsa05171). Shared AD-Covid mechanisms include: Nuclear Factor Kappa B (NFkB), Inhibitor of Nuclear Factor Kappa B Kinase (IKK), c-Jun N-terminal Kinase (JNK), Interleukin-6 (IL-6), Phosphoinositide 3-Kinase (PI3K), Tumor Necrosis Factor alpha (TNFa), TNF Receptor. Further, we found 22 genes involved in those AD-Covid common mechanisms and they correlate highly with AD phenotypes in different brain regions. Pathview^47^ was used to visualize the correlations of those genes and AD phenotypes. Also, for each brain region, we found the subnetworks of its GRN in AD that have TFs regulating those AD-Covid common genes as the region’s AD-Covid regulatory network.

### Gene expression analysis and machine learning prediction for Covid-19 severity from AD-Covid regulatory networks

To gauge the clinical predictive performance of our AD-Covid genes and networks in terms of predicting Covid-19 severities (and thereby immune system dysregulation), we looked at recent population RNA-seq gene expression data in blood of Covid-19 samples (GSE157103)^48^ to check whether any genes from our AD-Covid regulatory networks can predict Covid-19 severities (e.g. being in the Intensive Care Unit (ICU) or not). To this end, we median normalized this gene expression data (19,472 genes) and then applied differentially expression analysis by DESeq2^49^ between 50 ICU and 50 non-ICU Covid patients. Volcano plots highlighted differentially expressed AD-Covid genes (adjust p<0.05). A smoothing factor of 0.01 was added to the numerator and denominator when computing the empirical log2(fold change).

Apart from differentially expression analysis, which aims to find individual associated genes, we performed machine learning analysis for genes from our AD-Covid regulatory networks to see if they together can predict severe Covid-19 or not. In addition to our AD-Covid genes (from each region and combined), we also compared the machine learning prediction performance from other published Covid-19 genes. Since we predict Covid-19 severity, we compared predictive models from our lists with the respective performance of the benchmark list. A recent study^50^ using U.K. Biobank GWAS and Covid-19 mortality data discovered 8 Covid-19 susceptibility genes associated with an extremely high risk of Covid-19 mortality: DNAH7, CLUAP1, DES, SPEG, STXBP5, PCDH15, TOMM7, and WSB1. Another study^51^ identified 7 other risk genes (OAS1, OAS2, OAS3, TYK2, DPP9, IFNAR2, CCR2) associated with severe and life-threatening Covid-19 outcomes, including inflammatory organ damage. Numerous studies such as ^52^ implicated ACE2 and TMPRSS2 as key genes whose polymorphisms as risk factors for greater Covid-19 susceptibility. Another study using Random Forests machine learning to predict Covid-19 severity identified gene VEGF-D as the most important indicator for Covid-19 severity^53^. We thus included all 18 genes, from these published studies, into a list of Covid-19 published genes, to use as our benchmark list.

Python’s Scikit-Learn^54^ package, was used for our machine learning analysis. We used a Support Vector Machine (SVM) classification model (with linear kernel and balanced class weights, outputting predicted probabilities based on Python’s sklearn.svm.SVC package). The classification accuracy of each gene was calculated by stratified 4-Fold Cross Validation (CV); each fold had 25 unique human samples in the testing set (between 12 and 13 samples from each class) and the remaining 75 samples in the training set (between 37 and 38 samples from each class). For the 18 published Covid-19 susceptibility genes, we performed Recursive Feature Elimination (RFE) CV based on a SVM classification model; this calculated the classification accuracy of each gene and the optimal number of published genes to use (smallest number of genes with the maximum stratified 4-fold CV accuracy). We fixed all models to incorporate only this optimal number of genes. Then, to build a model for each of our input gene lists, we performed RFE based on our linear SVM to select this fixed optimal number of predictive genes from the list for classifying ICU vs. non-ICU Covid-19 patients with high accuracy (i.e., feature selection). We then trained an SVM classification model again with those select predictive genes and reported the average accuracy and AUROC values of the model using 4-Fold stratified CV. That is, we average the values for accuracy and AUROC across all 4 folds. Each of the 4 stratified CV folds trained a linear SVM kernel model, predicting the probability that a given Covid-19 positive sample would be in the ICU. Within each fold, we predicted these probabilities for our testing set of 25 samples, to get the predicted probabilities for our samples, which is input for our Decision Curve Analysis (DCA).

We used DCA^55^ to evaluate and compare the machine learning models of those brain-region AD-Covid genes and benchmark genes for predicting Covid severity. DCA is widely used for making medical decisions, improving upon traditional comparison metrics (e.g., AUROC) for predictive models and other approaches requiring additional information to address individuals’ clinical consequences for individuals^55^. Given a model and a threshold probability pT, patients will be sent to ICU if their percentage risks for Covid severity (i.e., ICU) from the model are greater than or equal to pT. Based on this, the true positive (TP) count is the number of Covid-19 severe individuals sent to the ICU, and the False Positive (FP) is the number of Covid-19 non-ICU individuals sent to the ICU. Thus, pT inherently represents subjective clinician preferences for FPs versus False Negatives (FNs: number of severe Covid-19 patients who were wrongly not sent to the ICU). Based on TP, FP and pT, the DCA then calculates Net Benefit = TP/N – ((FP/N)*pT/(1-pT)), where N is the total number of patients (N=100 here). Thus, the Net Benefit represents the benefit of true positive ratio (TP/N) from false positive ratio (FP/N) weighted by odds of pT (i.e., pT/(1-pT)). DCA provides a simple, personalized risk-tolerance based approach of using pT to weight the FN and FP mistakes: lower thresholds represent a fear of FNs over FPs, and vice-versa. For instance, for a clinician who sends a Covid-19 positive individual with predicted severity of at least pT = 20% to the ICU, the utility of treating a Covid-19 severe individual is 4 times greater than the harm of needlessly sending a non-severe Covid-19 patient to the ICU. We compared our predictive models with 2 extremes: Treat All (predict 1 for all Covid-19 positive patients and send all to ICU regardless of severity) and Treat None (predict 0 for all positive patients and send none to ICU). Practically, a clinician ought to opt for the predictive model or extreme intervention strategy with the highest Net Benefit based on that clinician’s preferred pT; thus, two clinicians (who may have their own, different pT values) may obtain different optimal results. Thus, DCA can evaluate the clinical usability of a Covid-19 severity prediction model based on its Net Benefit across clinically reasonable pT values. Finally, we performed DCA using code from Memorial Sloan Kettering Cancer Center^56^. Besides Covid severity, we also calculated gene expression correlations with Covid-19 and non-Covid for the genes from the AD KEGG pathway for three brain regions.

## Results

### Gene co-expression network analysis reveals gene expression dynamics for AD phenotypes across multiple brain regions

First, we applied our analysis to population gene expression datasets of three major brain regions relating to AD: Hippocampal CA1, Lateral Temporal Lobe (LTL), and Dorsolateral Prefrontal Cortex (DLPFC) (**Materials and Methods**). We identified several gene co-expression modules showing specific gene expression dynamic changes for various AD phenotypes as below. These expression dynamics also imply potential underlying gene regulatory mechanisms in the phenotypes. In particular, given a brain region, we constructed and clustered a gene co-expression network to a set of gene co-expression modules. In a gene co-expression network for a region, nodes are genes. Each edge represents that the 2 respective genes have correlated gene expression profiles during AD progression (i.e., co-expression). There are likely groups of co-expressed genes within the network form densely connected sub-networks (gene co-expression modules). Genes within a module share similar gene expression dynamics in the region for the observed AD phenotypes. We used modular eigengenes (MEs) to represent such expression dynamics for a gene module, using the first principal components of modular gene expression matrices. The information for all modules with associated phenotypes is available in **Supp. files 1, 2** and **3** for Hippocampus, LTL, and DLPFC, respectively.

#### Hippocampal CA1

We identified 30 gene co-expression modules in the Hippocampus. 21 modules were significantly positively associated with at least one phenotype: AD progression, Braak Stage progression, aging, accumulation of neurofibrillary tangles (NFTs), MMSE score, cognitive impairment, AD, and resilience. Also, their MEs show specific expression dynamics (**Fig. 2A** for 7 select modules, **Fig. S2** for all). For instance, pink and lightyellow modules have high expression values for Controls (no AD) and are clustered together. On the other hand, greenyellow, yellow, magenta, and tan modules have high expression levels for AD individuals and cluster together. In between those groups of modules is the midnightblue module, which has relatively high expression for some Control and some AD individuals.

**Fig 2.**
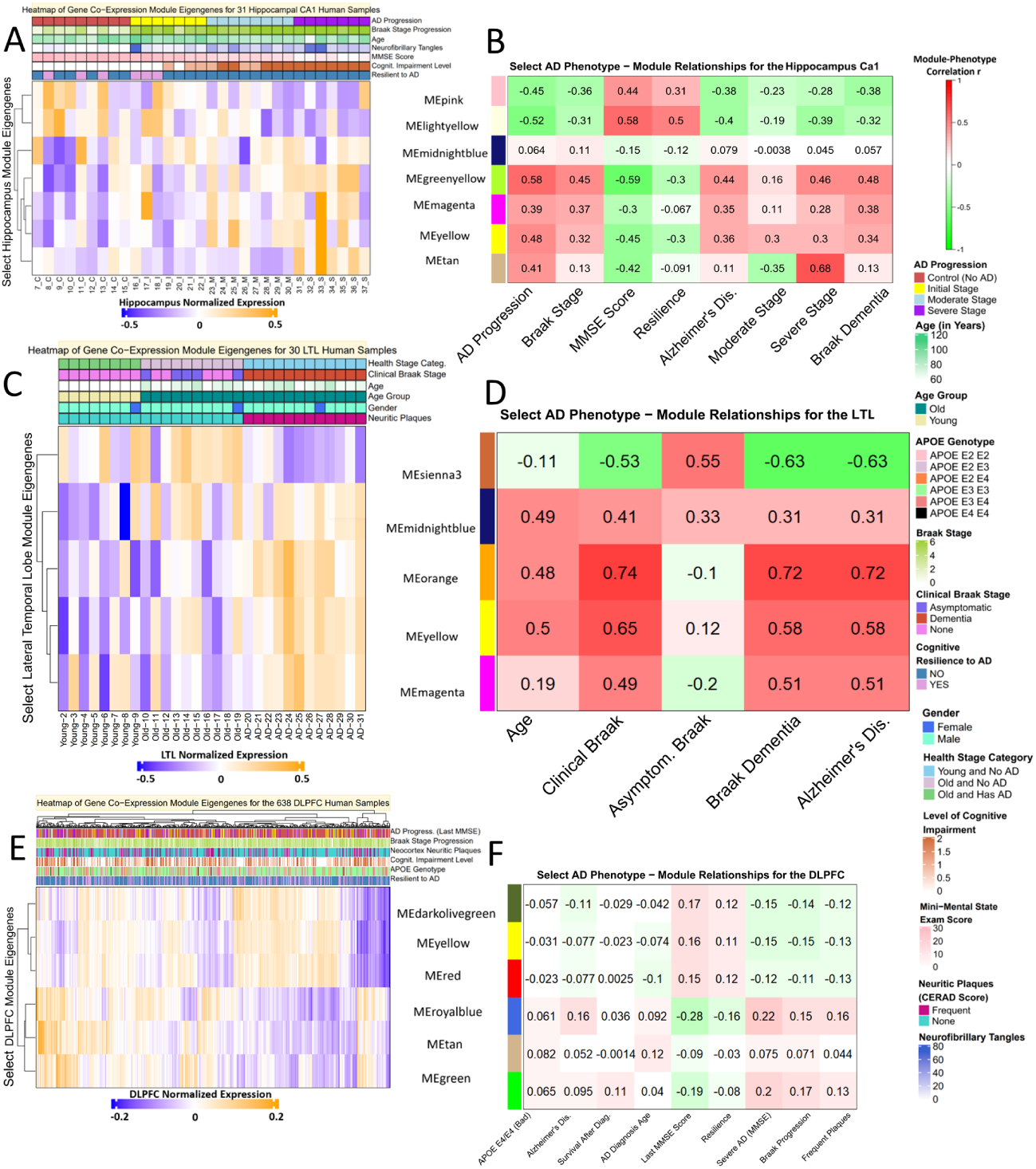
Gene co-expression modules significantly associated with AD phenotypes show specific expression dynamic patterns across phenotypes. Top heatmaps show the eigengenes of select gene co-expression modules in Hippocampal CA1 region (A), LTL (C) and DLPFC (E). Rows: modules. Columns: individual samples. Red: high expression level. Blue: low expression level. Bottom heatmaps show the correlation coefficients and p-values between select modules and AD phenotypes in Hippocampal CA1 region (B), LTL (D) and DLPFC (F). Row: modules. Columns: AD phenotypes. Red: highly positively correlation. Green: highly negatively correlation.

Next, using these expression dynamic patterns, we further linked these gene modules to key AD phenotypes (**Fig. 2B**) using their significant correlations (**Fig. S3** and **Supp. file 1** for all modules). For instance, the greenyellow module has significantly correlations with AD, AD progression, moderate and severe stages, Braak 6 stage and cognitive impairment. The tan module has the highest correlation (r = 0.68) with severe stage, along with other AD phenotypes. The midnightblue module is more significant (r = 0.41) for the Braak 4 stage, where affected individuals typically exhibit mild symptoms of dementia. The lightyellow module is significant for cognitive resilience (r = 0.5), which is the ability of individuals to exhibit stronger cognitive functioning despite AD pathology. Moreover, the lightyellow module strongly correlates with better MMSE performance (r = 0.58) than the pink module (r = 0.44). Instead, the pink module is more significantly correlated with the Braak 3 stage (r = 0.45).

#### Lateral Temporal Lobe

We identified 56 gene co-expression modules and found that 28 modules are significantly positively correlated with AD phenotypes of interest. We highlighted 5 MEs in **Fig 2C**, showing specific expression dynamics for AD phenotypes in LTL (**Fig. S4** for all modules). 12 gene modules are positively associated with AD Progression, 1 with Aging (mediumpurple3), 1 with Gender (lightgreen), 12 with Controls, and 2 with Initial Stage (associated with Braak 1 and 2 stages). As shown in **Fig. 2C**, the sienna3 module has higher expression values for both old and young individuals in Controls. Orange, magenta, and yellow modules are clustered together and have higher expression values for AD samples. The midnightblue module is clustered between both groups and tends to be associated with higher Braak stages. As shown in **Fig. 2D**, the sienna3 module also has a statistically significant positive correlation with Control Stage (r = 0.63) and asymptomatic based on the Braak stage (r = 0.55). The midnightblue module is associated with aging, average Braak stage, clinical Braak stage, and AD. The yellow, orange, and magenta modules are associated with aging, AD and Braak progression phenotypes, and neuritic plaque accumulation (based on CERAD score); the orange module has a very strong correlation with dementia Braak stages (r = 0.72) and AD (r = 0.72). More module-phenotype associations are available in **Fig. S5** and **Supp. File 2**.

#### Dorsolateral Prefrontal Cortex

We found 35 gene co-expression modules for DLPFC. **Figs. 2E** and **2F** highlighted 6 of those gene modules: darkolivegreen, yellow, red, royalblue, tan, and green (**Figs. S6-7** for all modules). Those modules are significantly associated with various AD phenotypes such as progression, MMSE, APOE genotype, neuritic plaques. Larger sample size for the DLPFC (over 20 times larger than that for Hippocampal or LTL), likely attributes to relatively lower correlation coefficients between modules and AD phenotypes in DLPFC. However, we still see significantly correlated modules with various AD phenotypes (**Fig. 2F**, p< 0.05). For example, the tan gene module is associated with the worst APOE genotype (E4/E4) (r = 0.082), AD diagnosis age (r = 0.12). The royalblue and green modules are statistically significantly positively correlated with severe stage based on last MMSE score, with correlations of r = 0.22 and r = 0.2, respectively. In terms of better outcomes, the darkolivegreen module is significant for Controls (r = 0.11), better performance on last MMSE (r = 0.17), and cognitive resilience (r = 0.12). Furthermore, the red module is significant for better performance on last MMSE (r = 0.15) and has a similar correlation as darkolivegreen module for cognitive resilience.

Therefore, those AD-phenotype-associated gene co-expression modules uncover the specific gene expression dynamic patterns across phenotypes and suggest that those co-expressed genes in the same modules are likely involved in similar functions and pathways. To understand this, we performed enrichment analysis of the modules as follows.

### Eigengenes and enrichments of co-expression modules reveal hub genes, gene functions, and pathways in AD phenotypes

Gene module enrichment analysis allows us to better understand biological functions, structures, diseases, and other observed biological phenomena associated with AD phenotypes. Through enrichment analysis (**Methods**), we found enriched functions and pathways of AD modules and linked them to various AD phenotypes associated with the modules (**Fig. 3** and **Supp. document**). Overall, healthier phenotypes are Control Stage (no AD), cognitive resilience, and protective APOE E2/E2 genotype.

**Fig 3.**
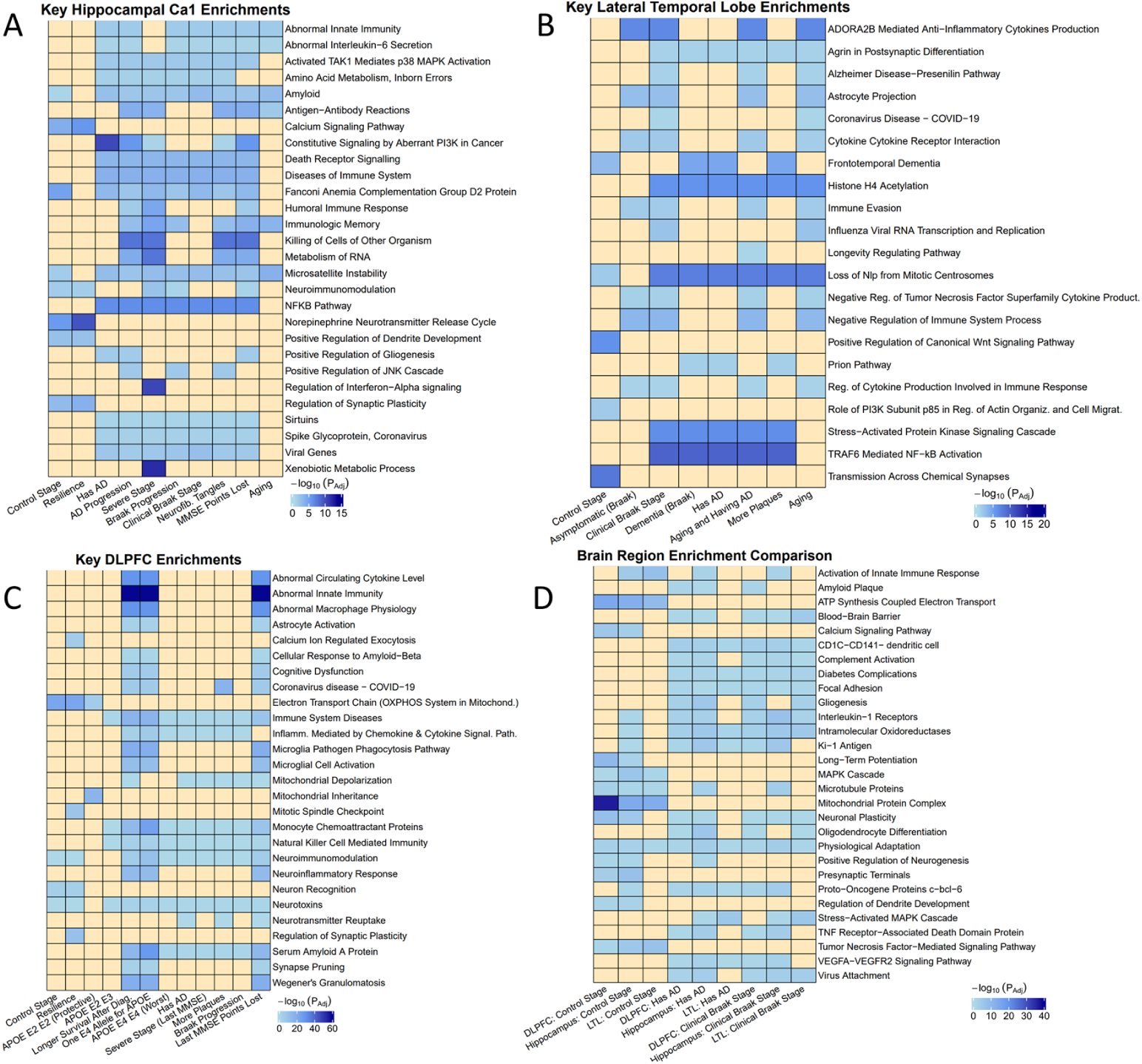
Select enriched functions and pathways of gene co-expression modules for various AD phenotypes. (**A**) Hippocampus; (**B**) Lateral Temporal Lobe (LTL); (**C**) DLPFC; (**D**) Across three regions. Rows: select enriched terms (**Methods**). Columns: AD phenotypes. The heatmap colors correspond to –log10(adjust p-value).

#### Hippocampal CA1 (Fig. 3A, Supp. file 1)

The Hippocampus CA1 (crucial for autobiographical memory, mental time travel, self-awareness) has a significant loss in neurogenesis, memory ability, volume, and neuronal density in AD^57^. Modules for non-AD phenotypes are enriched with synaptic plasticity and dendrite development, norepinephrine neurotransmitter release cycle, and calcium signaling pathway, which can all be dysregulated in AD. This also suggests that resilient individuals may be protected from microsatellite instability and amyloid accumulation that may even occur naturally. Further, our analysis supports recent hypotheses on dysregulated immune systems in AD. Aging, NFTs, and AD developmental phenotypes are associated with abnormal innate immunity and IL-6 secretion. Our cognitive impairment, Braak progression, and AD modules are enriched for: viral genes, Covid-19 spike glycoprotein, activated TAK1 mediating p38 MAPK activation (linked to tau phosphorylation, neurotoxicity, neuroinflammation, synaptic dysfunction, worse AD^58^), NFKB pathway (impaired and over-expressed during AD, leading to neuroinflammation, microgliosis, suppression of Wnt Signaling^10^). We found an association between Severe AD and immunologic memory, antigen-antibody interactions, and regulation of Interferon-Alpha Signaling. Interferon (cytokines mediating host response to viral infection with high expression in Hippocampal tissues of AD mice^59^) response to immunogenic amyloid may activate microglia, initiate neuroinflammation, and lead to synaptic loss^60^. Many AD phenotypes are associated with Death Receptor Signaling, positive regulation of gliogenesis, Constitutive Signaling by aberrant PI3K in Cancer, positive regulation of JNK cascade (activated in AD brains, involved in tau phosphorylation and neuronal death)^70^. Xenobiotic Metabolic Processes are specific to Severe AD modules only, and studies^61^ found links between dementia progression and various metabolic pathways.

#### LTL (Fig. 3B, Supp. file 2)

The LTL contains the cerebral cortex (responsible for hearing, understanding language, visual processing, facial recognition) and is impacted early in AD^25^. Control modules are enriched with several pathways typically present in healthy conditions like Wnt signaling, Actin organization, and transmission across chemical synapses. Dysregulation of those pathways can lead to neurodegeneration, e.g., Wnt signaling to inhibit amyloid-beta production and tau protein hyperphosphorylation in AD progression^62^. Second, in LTL modules for AD and neuritic plaques, loss of Nlp from Mitotic centrosomes is enriched (may lead to reduced microtubule stability, abnormal cellular morphology, and functions in AD^63^). Also, we found cell-type associated enrichments in the AD progression-related phenotypes, e.g., astrocyte projection for clinical Braak stage and asymptomatic. Astrocytes are increasingly activated near amyloid plaques in LOAD, producing pro-inflammatory cytokines and reactive oxygen species^64^. Other AD-related functions and pathways include postsynaptic differentiation, stress-activated signaling, TRAF-mediated NF-kB activation, prion pathway, epigenetic modifications. The prion pathway feedback loop is likely disrupted during AD leading to Aβ accumulation^65^. Epigenetic modifications (e.g., Histone A4 acetylation) can be dysregulated to affect gene expression of long-term potentiation and memory formation in AD; a recent study observed dramatic H4K16ac losses near genes linked to aging and AD in LTL^25^. AD-related phenotype modules are more enriched for Frontotemporal Dementia than controls are.

#### DLPFC (Fig. 3C, Supp. file 3)

The DLPFC is involved in executive functioning, supports cognitive responses to sensory information, works with the Hippocampus to mediate complex cognitive functions^66^, has plasticity deficits in AD^67^. Microglia exclusively express many AD risk genes like APOE^68^. Our APOE E2 modules are shielded from neurotoxins and associated with mitochondrial inheritance (p<1e-16), while E4 modules are strongly enriched for cellular response to Aβ and cognitive dysfunction. Having both high-risk E4 alleles (instead of 1) is not associated with mitochondrial depolarization but more strongly associated with Serum Amyloid A Protein (p<1e-28 vs. p<1e-16), neuroimmunomodulation, monocyte chemoattractant proteins.

We found several strong promising associations (some with p<e-58) for APOE4-related and cognitive impairment modules, supporting the crucial role of neuroimmunology and reactive microglia in AD pathogenesis. APOE4 signaling may regulate microglial response to Aβ^88^; our enrichments may help shed light on links between APOE4 and AD neuroinflammation, which is still unclear. These include astrocyte activation (boost production of proinflammatory cytokines and phagocytic capabilities), abnormal circulating cytokine levels and innate immunity, synapse pruning, neuroinflammatory responses, autoimmune diseases (Wegener’s Granulomatosis), abnormal macrophage physiology, Microglia Pathogen Phagocytosis Pathway, microglial cell activation. During the early synaptic decline in AD, microglia may change shape, functions, and pathways, express more receptors, become more phagocytic and activated, and go awry, releasing numerous pro-inflammatory cytokines, leading to chronic neuroinflammation, cell death^68^, neurodegeneration, and worse Aβ and NFT pathologies^8^. The prefrontal cortex of AD brains (along with the Hippocampus) has increased proinflammatory cytokines like IL-1B, associated with Aβ plaque deposition and higher AD risk^8^. Except for neurotransmitter reuptake, other AD phenotypes (ex. severe stage, Braak progression, plaques, cognitive impairment, survival post-diagnosis) modules share many biological associations with APOE4 modules, like Immune System diseases, Covid-19, inflammation mediated by chemokines and cytokines, natural killer mediated immunity. Non-AD phenotype modules are enriched with dysfunctional pathways in AD such as Electron Transport Chain^69^, neuron recognition, calcium ion regulated exocytosis, mitotic spindle checkpoint, and synaptic plasticity.

#### Comparison Across Brain Regions (Fig. 3D)

Many enriched pathways for AD phenotypes are shared by different brain regions. In particular, the datasets of 3 brain regions share major phenotypes (control, AD, clinical Braak stage) enriched with physiological adaptation. Control and AD Hippocampal and DLPFC modules share neuroimmunomodulation. The MAPK cascade (associated with control modules in all three regions) is associated with stress-activation in AD and Braak modules in Hippocampus and LTL. AD and Braak modules in multiple brain regions are also enriched with Ki-1 antigen (tumor marker of activated immune cells regulating NF-kB and apoptosis), proto-oncogenes, dendritic antigen-presenting cells, diabetes complications, focal adhesion (plaques), angiogenesis VEGFA-VEGFR2 Signaling. VEGFA-VEGFR2 pathway promotes neural cell survival, migration, and proliferation and has altered levels in AD that may impact microglia^70^. Moreover, AD and Braak modules across regions have common enrichments such as Blood-Brain Barrier, virus attachment, Complement Activation (innate immune-mediated defense altered in AD^71^), and Oligodendrocyte differentiation (potentially associated with Aβ accumulation and neurodegeneration^72^). Finally, control modules across regions are enriched for higher cellular energy levels like ATP synthesis, Mitochondrial proteins. DLPFC and Hippocampal control modules are enriched with Calcium Signaling, LTP, Presynaptic Terminals, regulation of dendrite development, positive regulation of neurogenesis.

Overall, our module enrichments across brain regions further underscore the role of neuroimmunology and other factors in AD progression across the regions and verify those phenotype correlations we detect for our gene co-expression modules are potentially indicative of true biological signals. In **Figures S8-S10**, we identify select gene co-expression modules with strong biological enrichments and significant correlations with AD phenotypes.

### Prediction of brain-region gene regulatory networks for AD phenotypes

To understand underlying molecular mechanisms regulating gene expression associated with various AD phenotypes, we predicted the gene regulatory networks (GRNs) for genes and gene modules of brain regions, especially using multi-omics data (**Methods**). The brain-region GRNs link transcription factors (TFs) and regulatory elements (REs, e.g., enhancers or promoters) to target genes (TGs) and co-expressed genes (e.g., from the same module). Regulatory network edges can be activation or repression, which follow-ups can investigate. Moreover, these GRNs can be further linked to the AD phenotypes significantly associated with TGs and modules. As described in **Methods**, we applied multiple widely-used approaches and public databases to predict networks and used shared predictions across different approaches as highly confident GRNs. In terms of candidate TFs, we found: 1,043 in Hippocampus, 1,580 in LTL, and 1,588 in DLPFC, which we input into RTN, Genie3, and Trena. As shown in **Table S2**, we obtained 6,823,631 TF-RegulatoryElement-TG network edges of 3 brain regions’ GRNs, corresponding to 973,025 unique TF-TG pairs, 20,601 TGs, and 709 TFs. In particular, the Hippocampal GRN has 2,810,102 TF-RegulatoryElement-TG edges, including 169,292 unique TF-TG pairs, 11,972 TGs and 351 TFs. The LTL GRN has 161,404 TF-RegulatoryElement-TG edges, including 65,321 unique TF-TG pairs, 13,791 TGs and 402 TFs. The DLPFC GRN has 3,852,125 TF-RegulatoryElement-TG edges, including 752,169 unique TF-TG pairs, 13,511 TGs and 670 TFs. Detailed edge lists of Hippocampus and LTL GRNs are in **Supp. Files 4-5**.

### Identification of disease risk variants for AD phenotypes via integration of GWAS and gene regulatory networks

Over 90% of disease risk variants (e.g., GWAS SNPs) are in non-coding regions^73^. We found that GWAS SNPs for AD are enriched in the regulatory elements of our GRNs as above. Thus, it is crucial to further understand how those disease risk variants affect gene regulatory mechanisms that eventually impact AD phenotypes. To this end, we linked AD GWAS SNPs to our GRNs to see how those SNPs interrupt TFBSs on enhancers or promoters that regulate target genes and modules (**Methods**). These SNPs can be linked to various AD phenotypes of corresponding genes and modules for different brain regions, i.e., “brain-region risk variants for AD phenotypes.” Specifically, 39,832 unique AD SNPs disrupted TFBSs on regulatory elements of 3 brain-region GRNs (35,940 for Hippocampus, 7,119 for LTL, 2,359 for DLPFC, **Fig. S11**). Across three regions, there are 543 unique TFs whose binding sites were interrupted, regulating 11,596 genes (**Table S3**). Our analysis links many non-coding disease-associated SNPs to different phenotypes, and here we highlight a select few different examples for key AD genes.

For instance, a subnetwork of the LTL GRN between TFs is shown in **Fig. 4A**, i.e., the TGs are also TFs. These subnetwork TFs also have interrupted TFBSs on the regulatory elements to their target genes. Several TFs are hub genes of the subnetwork like RUNX2 (11 TFs have difficulty binding and regulating RUNX2) and neurogenesis TF REST (experiences difficulty in regulating 16 TFs in LTL). REST is induced by Wnt signaling, represses genes (like PLCG2) that promote cell death or AD pathology, protects neurons from Aβ-protein toxicity^74^. We found that during AD, PLCG2 is overexpressed in the LTL, which may be partly explained by REST’s inability to bind to chromatin and repress its target genes^75^; this may lead to changes in autoinflammation, immune disorders, immune cell functioning^76^. Moreover, REST significantly regulates a gene module of 883 genes (turquoise; hub gene: BOLA2-SMG1P6) in LTL (**Figs. S12-13**). The subnetwork of DLPFC GRN between TFs (**Fig. S14**) has hub genes: CREB3L1 (26 TFs are unable to properly regulate CREB3L1) and PAX5 (difficulty in regulating 11 TFs). Lastly, the Hippocampal GRN SNP subnetwork between TFs has hubs such as ZNF226 (21 TFs experience difficulty regulating ZNF226) and GATA2 (unable to regulate 65 TFs in AD Hippocampus). In the Hippocampus, ZNF226 significantly regulates three modules (2 which are associated with Controls), and GATA2 significantly regulates 4 modules (1 Control Stage module and 2 modules associated with worsening AD phenotypes) (**Figs. S15-16**).

**Fig 4.**
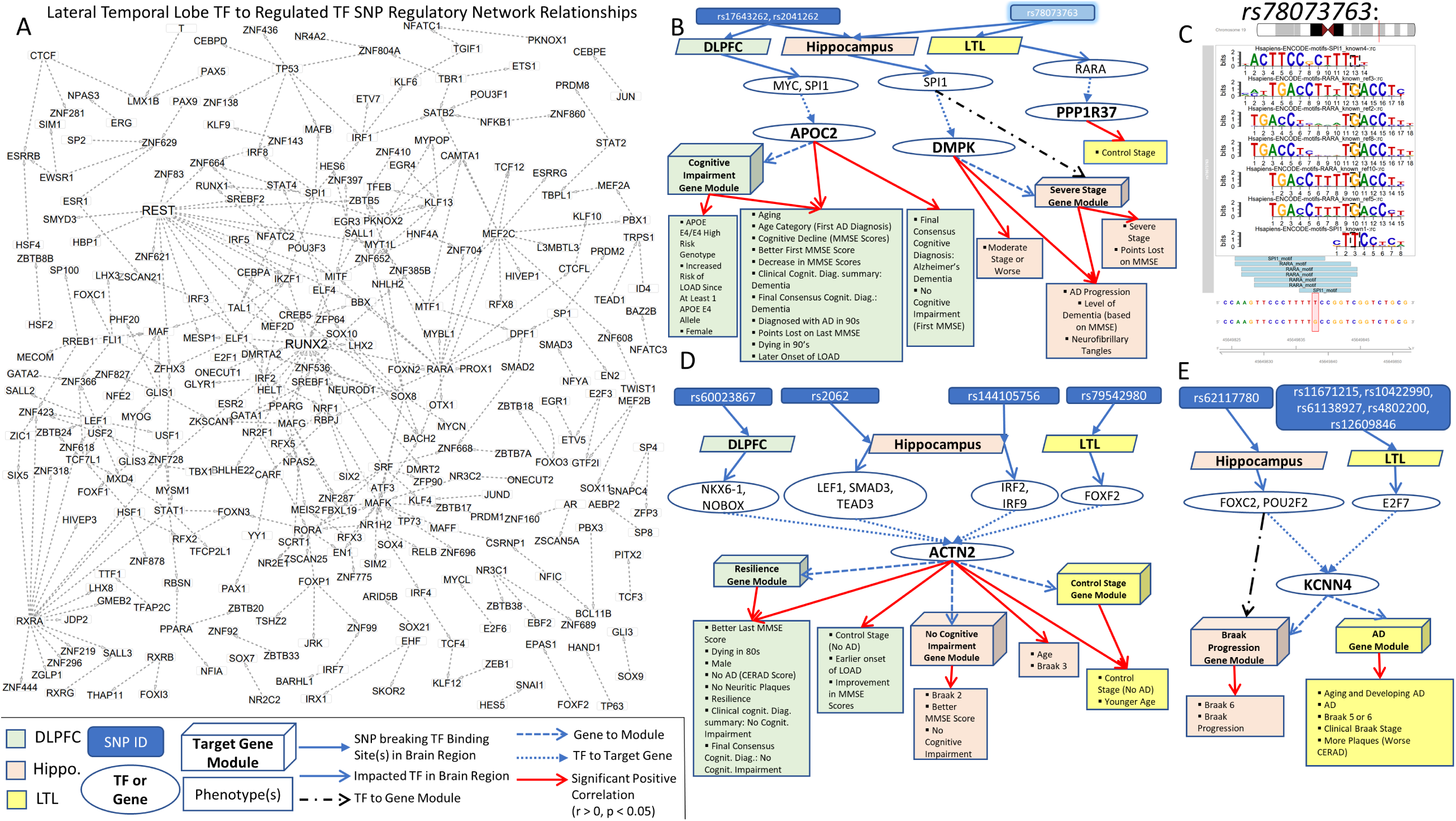
Select gene regulatory networks (GRN) linking AD risk variants (GWAS SNPs) to AD phenotypes. (**A**) Subnetwork of LTL GRN among TFs (i.e., the target genes (TGs) are TFs too). Nodes are TF genes. The edges connect TFs to their TGs. Besides, TFs have the binding sites interrupted by AD SNPs on the regulatory elements to TGs. (**B**) Example of 3 AD SNPs that interrupt binding sites of TFs in different brain regions. The AD phenotypes and gene modules positively correlated with APOC2, DMPK, and PPP1R37 expression are shown. (**C**) SNP rs78073763 interrupts multiple possible binding sites of SPI1 in Hippocampus and RARA in LTL. Additional regulatory links from AD SNPs to interrupted TFs to TGs along with associated phenotypes and modules are shown in (**D**) regulation of ACTN2 in all three regions, (**E**) FOXC2, POU2F2, and E2F7 to KCNN4 in Hippocampus and LTL.

Furthermore, we found several regulatory networks, which provide more insights on the possible association of various non-coding SNPs with AD phenotypes. **Fig. S17** provides a detailed explanation of how to interpret such networks. In **Fig. 4B**, we examine the varying effect of a given SNP on gene regulation across brain regions and the impact on 3 TGs: APOC2, DMPK, and PP1R37. SNP rs78073763 (**Fig. 4C**) changes the DNA base from a T to a G (at chr19:45649838) and breaks the binding of RARA on the enhancer of Control gene PPP1R37 in LTL and SPI1 binding to DMPK Hippocampus enhancer. A recent study found PPP1R37 expression is strongly associated with APOE expression and has extensive cross-tissue effects on AD, and DMPK’s Hippocampal expression strongly impacts AD^77^. We found increased DMPK’s Hippocampal expression is associated with worsening AD phenotypes (ex. Moderate stage or worse, AD progression, more severe dementia). 2 extremely statistically significant AD SNPs (rs17643262 and rs2041262; p < 2e-13) disrupting SPI1 regulation of DMPK in the Hippocampus also disrupt SPI1 regulation of APOC2 in DLPFC; APOC2 is associated with a cognitive impairment tan DLPFC gene module, AD dementia, cognitive decline, having at least 1 APOE E4 allele. APOC2’s tan DLPFC gene module is significantly associated with many meaningful biological enrichments, including Covid-19, neuron death, neuroinflammatory response, abnormal innate immunity, TNFa signaling via NFKB, response to Interferon Gamma, brain death (**Fig. S10**). SPI1 is a well-known master regulator in microglial cells, plays a key roles in regulating immune functions in AD^78^, is strongly correlated with AD (r = 0.355) and AD Progression (r = 0.375), Braak progression (r = 0.437), and Braak 6 (r = 0.407), and belongs to a severe stage gene module (r = 0.41). This suggests low-level SPI1 expression in Hippocampus controls, potentially reducing microglial-mediated neuroinflammatory responses and delayed AD onset^79^. SPI1 regulation in the Hippocampus by 10 TFs (ex. RXRA, RARA, NFKB1) is disrupted by several SNPs (**Fig. S18**). SPI1’s regulated genes are also upregulated in microglia, leading to microglia-mediated neurodegeneration in AD^79^; in fact, SPI1 significantly regulates DMPK and its Severe Stage Hippocampal gene module. MYC regulation of APOC2 in DLPFC is also impacted, possibly leading to dysregulation in cellular processes like cell cycle activation and Wnt Signaling and re-entry mediated neuronal cell death in AD^80^.

Since non-coding SNPs may have highly cell-type specific effects, we investigated the epigenetic landscape and regulatory element signals for these three significant putative functional SNPs (rs78073763, rs17643262, rs2041262) that are very close to PP1R37 (co-expressed with APOE) and also impact binding of SPI1 (**Fig. S19**). Microglia-specific regulatory elements are indeed present in these SNPs, impacting SPI1 binding and regulation of microglia. Our results further underscore dysregulated microglia and neuroimmunology in AD.

In **Fig. 4D**, we present an example of a gene, ACTN2 (cytoskeletal alpha-actin protein), whose regulation in all three brain regions are impacted by various SNPs and whose expression is associated with healthier outcomes. Across the three brain regions, we observe that ACTN2 expression is positively associated with cognitive resilience DLPFC gene module, no cognitive impairment Hippocampus module, and Control LTL module. Increased ACTN2 expression may be associated with better MMSE scores; ACTN2 is found at the neuronal synapse and is highly ranked in a previous study on echocardiographic traits, heart function, and AD^81^. Several SNPs impact ACTN2 DLPFC and Hippocampal expression, but we focused on SNPs shared by several TFs in this Figure. Here, rs60023867 disrupts the regulation of ACTN2 in DLPFC by both NKX6-1 and NOBOX. In the Hippocampus, rs2062 disrupts the regulation of ACTN2 by 3 TFs (LEF1, SMAD3, TEAD3), and rs144105756 impacts IRF2 and IRF9 regulation of ACTN2. Lastly, in the LTL, rs79542980 may lead to FOXF2 dysregulation of ACTN2.

Different Hippocampal and LTL SNPs impact regulation of KCNN4, a key AD drug target overexpressed during AD (**Fig. 4F**). In **Fig. S20**, we visualize the impact of rs62117780 on FOXC2 and POU2F2 regulation of KCNN4 in the Hippocampus and rs4802200 on E2F7 regulation of KCNN4 in the LTL. FOXC2 and POU2F2 also regulate KCNN4’s Hippocampal Braak progression module. KCNN4 belongs to the magenta AD LTL module, so increased KCNN4 expression is associated with AD progression in both regions. Furthermore, KCNN4’s LTL module has key biological enrichments, like TNFa/NFkB Signaling Complex, death receptor signaling, autoinflammatory disorder (**Fig. S9**). SNP rs62117780 is in a microglia signal peak (**Fig. S21**), consistent with previous findings that KCNN4 is mainly expressed in microglia and regulates microglial activation by modulating Ca2+ influx signaling and membrane potential^75^. KCNN4 has low expression in healthy neurons and is associated with neuroinflammation and reactive gliosis during AD. Blocking KCNN4 likely curbs microglial neurotoxicity, leading to slower neuronal loss and better memory levels^82^. Therefore, this link uncovers how AD SNPs regulate KCNN4 expression in AD phenotypes.

Finally, we highlighted all possible SNPs that interrupt TFBSs in our brain region GRNs via Manhattan plots (**Figs. S22-25**): **S23**(Hippocampus), **S24** (LTL), **S25** (DLPFC). 268 SNPs were found in all three regions and are examined in **Figs. S26-29**. Regulatory links from AD SNPs to interrupted TFBSs and regulatory elements to target genes and modules are provided in **Supp. Files 6-8**. We provide additional examples and explanations of regulatory networks linking non-coding SNPs to AD phenotypes for the networks, which we visualize in **Figs. S30-33**.

### Gene regulatory networks and AD phenotypes associated with shared pathways between Covid-19 and AD

To better understand the role of the dysregulated immune system in AD onset and progression, we utilized the recent Covid-19 virus that has widely affected the elders with neurogenerative diseases, including AD, since there are potential links between Covid-19 and AD. As rogue immune responses characterize both diseases, we look at shared AD-Covid pathways to understand molecular mechanisms that may be implicated in adverse effects and inflammation in both diseases. One such pathway is the NFKB pathway.

Five TFs: NFKB1, NFKB2, REL, RELA, and RELB (proto-oncogene near APOE) are involved in this pathway. In AD and Covid-19, Reactive Oxygen Species (ROS) activate RELA and NFKB1 that then transcribe pro-inflammatory cytokines (typically secreted by macrophages), like IL-6, IL-1B, and TNF, reducing LTP in AD, leading to an exaggerated and potentially lethal immune response in Covid-19 (e.g., tissue injury, Acute Respiratory Distress Syndrome^83^, hypoxia^13^) (**Fig. S34**). In general, NFKB gene expression levels positively correlate with AD phenotypes such as severity but negatively with control in all three regions (**Fig. 5A** for Hippocampus). For instance, NFKB1 and RELB negatively correlate with controls in 3 regions, and so do NFKB2 and RELA in Hippocampus and DLPFC (**Fig. 5B**). However, all 5 TFs positively correlate with AD severity in the Hippocampus and 2 of them in DLPFC (**Fig. 5C**), which may help support current AD hypotheses that NFKB activation and cellular inflammation is a primary cause of AD^10^. Since up-regulation and activation of NF-kB TFs are linked to greater inflammatory responses in Covid-19 and AD^83^, NFKB gene expression may be a potential interplay between AD and Covid-19. We further investigated our GRNs involving NF-kB TFs to understand possible regulatory mechanistic links across AD and Covid-19 and how NFKB dysregulation may be linked with AD neuroinflammation. Improper NFKB functioning may be associated with improper immune functioning, altered neuronal dynamics (pushing neurons towards degeneration), microglial activation, oxidative stress-related complications, over-production of Aβ, and excessive inflammation, observed in memory disorders such as AD^10^.

**Fig 5.**
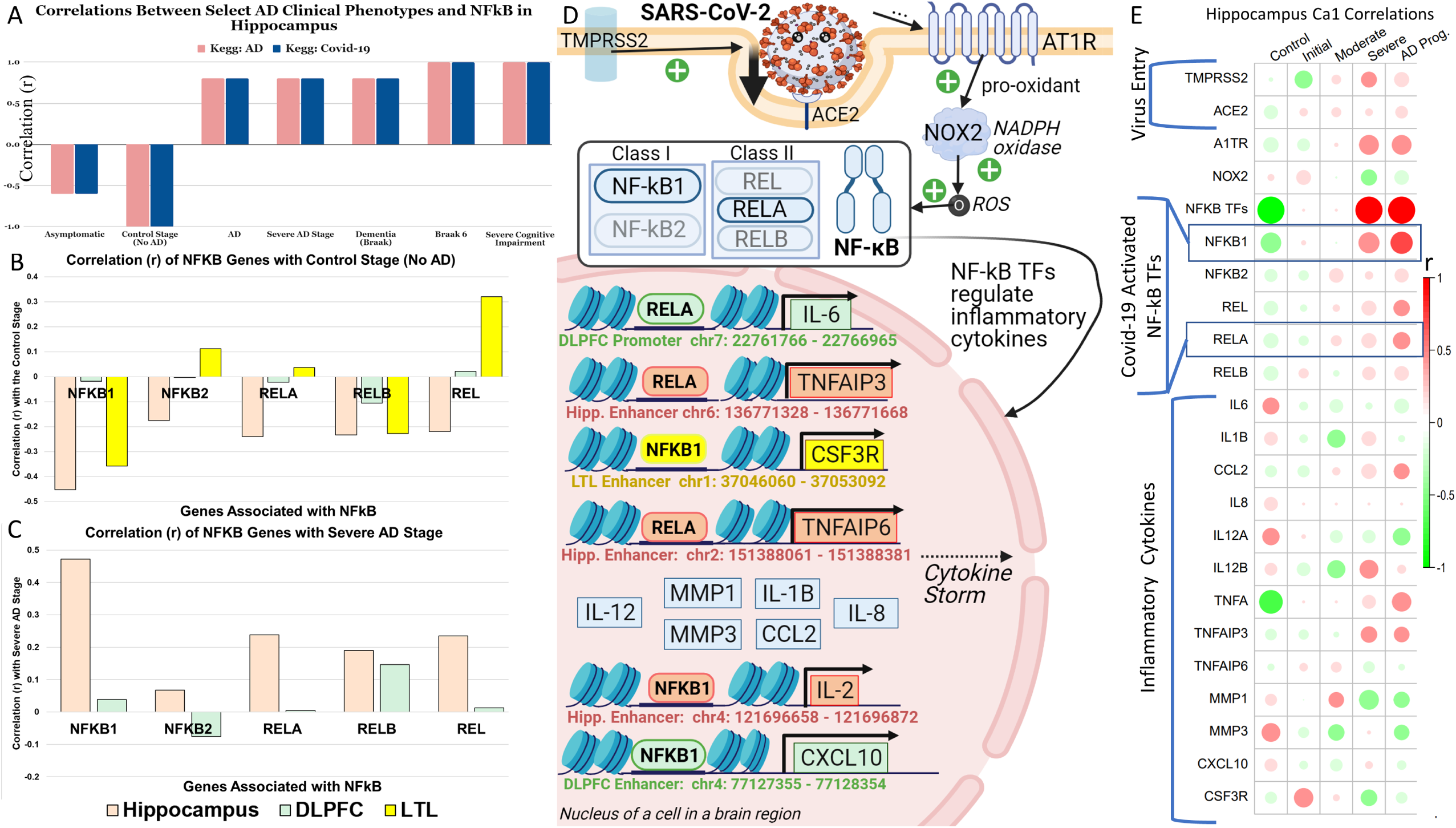
Gene regulatory networks and phenotypes for NFKB, a shared pathway of AD and Covid-19. (**A**) Correlations of the NFKB pathway (KEGG:hsa05171) and AD pathway (KEGG: hsa05010) with AD phenotypes from the Pathview analysis of Hippocampal expression data of pathway genes^47^. (**B**) Correlations of NFKB TFs (NFKB1, NFKB2, RELA, RELB, and REL) with Control in three regions. (**C**) Correlations of NFKB TFs with Severe stage in Hippocampus and DLPFC. (**D**) Covid-19 virus spike and gene regulation of NFKB TFs from our Hippocampal, LTL, and DLPFC GRNs for pro-inflammatory cytokines linked with severe Covid-19 outcomes. Gray dashed arrows indicate regulation and black arrows indicate activation of cytokines by the TF. (**E**) Correlations between AD phenotypes and expression levels of genes from (**D**) in the Hippocampus.

To this end, we looked at the impact of the NFKB pathway in SAR-CoV-2 on additional molecules (KEGG: hsa05171) (**Fig. 5D**). In particular, to enter the cell, the SARS-CoV-2 Spike protein is primed by TMPRSS2, binds to ACE2 (highly expressed in macrophages and the brain^84^), and interacts with AT1R (elevates viral entry and infection^85^). Neural cells may be directly invaded by SARS-CoV-2 or by systemic infection that compromises the Blood Brain Barrier (is even more dysfunctional in AD), elevating levels of cytokines and chemokines in the brain that encounter astrocytes and microglia (that are also dysfunction in AD)^86^. Expression levels of TMPRSS2, ACE2, and AT1R receptors are negatively correlated with controls but positively associated with late AD stages in Hippocampus (**Fig. 5E**).

Moreover, in our brain region GRNs, NFKB1 and RELA regulate genes of several cytokines associated with the severe Covid-19 Cytokine Storm. For instance, in the DLPFC, RELA regulates IL-6 via the promoter and particularly activates CXCL10 (biomarker whose altered levels are associated with immune dysfunction, tumor development, and disease severity^87^) by binding to the enhancer. In the LTL, NFKB1 binds to the enhancer of cytokine CSF3R, a key regulator of neutrophil development, proliferation, and differentiation, essential for development and maintenance of microglia^88^; in Severe Covid-19 patients, there are increased levels of neutrophils (immune cells involved in first-line of defense against pathogens) and changes in their phenotype and functionality^89^. CSF3R mutations are associated with brain structure abnormalities and our pipeline has identified several non-coding SNPs that impact CSF3R regulation by STAT3, TP53, EGR2, and IRF8 in Hippocampus by SPI1, REST, NFKB1, and NFATC2 in LTL. For instance, rs12120626 (p = 0.000319199) disrupts STAT3 regulation of CSF3R in Hippocampus and SPI1 regulation of CSF3R in LTL; rs6425995 (p = 0.000818697) disrupts CSF3R regulation by REST and NFKB1 in the LTL.

Also, our Hippocampal GRN found that NFKB1 regulates IL-2 (via an enhancer on chr4:121696658-121696872) and RELA regulates TNFa-induced proteins (regulate long-term potentiation), TNFAIP3 and TNFAIP6. Overall, NF-kBs regulate TNF-a, increasing expression during AD progression, likely triggering neurodegeneration, inflammation, neuronal death, healthy tissue destruction, increasing Aβ burden and initiating a cytokine cascade^8^; research suggests that Aβ stimulation may activate NFKBs that upregulate the expression of pro-inflammatory cytokines like TNFa and IL-1B in microglia and astrocytes^10^, which then increase Aβ burden and plaque deposition in AD brains^8^.

We analyze how specific genes involved in the NFKB pathway in Covid-19 correlate with AD phenotypes to help us better understand key AD genes (targeted in Covid-19) that may drive neuroinflammation in AD. RELA regulates genes of several inflammatory cytokines associated with the severe Covid-19 Cytokine Storm, and studies suggest that RELA may be the most important TF regulating Covid-19 response^91^. The Covid-19 cytokine storm has been associated with BBB dysfunction, cellular senescence, astrocyte injury and anti-neuronal antibodies, and activation of microglia and astrocytes, causing neuroinflammation, neuronal death, widespread neurodegeneration^13^. We found that RELA regulates IL-6 and particularly activates IL-12A/B (recruit and activate Natural Killer cells) and IL-1B via enhancers. In **Fig. S32**, we shared additional examples of NFKB1 and RELA TFs regulating other TFs that regulate inflammatory cytokines IL-1B (regulates APP synthesis), IL-12B, CCL2, MMP 1/3, and CLGN. In the Hippocampus, NFKB1 regulates TFs SPI1 and BATF, regulating MMP1 (**Fig. S32B**). Expression of cytokines IL-2, CCL2, IL-1B IL-12B, and TNFa highly positively correlate with AD severity (**Fig. 5E**). IL-2 and TNFa are usually highly expressed in Covid-19 patients with severe pneumonia who develop ARDS and need intensive care and oxygen therapy^83^. NFKB1 and RELA belong to the same AD progression and Severe AD Stage greenyellow Hippocampal gene module of 383 genes, which has many immunological enrichments, such as: microglial cells, Interleukin-1 receptors, death receptor signaling, PID IL-1 Pathway, TLR2 Signaling, abnormal innate immunity and immunoglobulin level, positive regulation of innate immune system response, and NFKB is activated and signals survival (**Fig. S8**).

Further, APOE genotype is associated with differences in Complement Cascade Component C1qrs expression in Covid-19 patients^15^ in the DLPFC, as it is negatively correlated with E2/E2 but positively with E4/E4 (**Fig. S36**). C1qrs activates microglia to the M1 state, releasing mediators that increase inflammation and damage healthy cells^93^. We found that C1qrs is negatively correlated with Control and Initial Stages, but positively correlated with Moderate and Severe Stages in the Hippocampus (**Fig. S37-38**). Complement activation is involved in an inflammatory feedback loop with neutrophil activation (resulting in tissue injury)^94^, and may be a hallmark of severe Covid-19. Many complement components are negatively correlated with Control and Initial Stages in the Hippocampus (except MBL and VWF), but positively correlated with AD progression. IgG antibodies, whose responses to various epitopes are key to the immune response to Covid-19^95^, are the only component positively associated with Moderate AD but not severe AD. Fibrinogen and SELP changed from negative to positive associations from Moderate to Severe AD stages.

Additionally, our GRNs help find potential roles of AD SNPs in dysregulating expression of NFKB TFs, which will have reverberations in not only Covid-19 but also AD. In **Fig. S32**, we examine the impact of various AD SNPs on the expression of NF-kB TFs and NFKB regulation of key cytokines described in **Fig. 5D**. NFKB TFs may be activated in several different ways, including canonical and non-canonical pathways. We found several SNPs disrupting regulation of RELA (by 5 TFs) and NFKB1 (by 7 TFs including SPI1 and REST) in the Hippocampus (**Fig. S39**). This may help us understand the mechanisms of NFKB TF activation in AD and identify specific activators and is important because: NFKB TFs may be either neuroprotective or neurotoxic based on specific activators^96^ and dominant activators of NFKBs in AD are still unknown. Moreover, there are 4 highly significant SNPs (p < 5e-15) impacting RELA’s ability to regulate ZNF226, a hub TF from the Hippocampus SNP TF-TF Subnetwork GRN (**Fig. S32B**); RELA also regulates ZNF226’s gene module. 99 extremely significant SNPs (p < 1e-9) impact regulation of RELB in DLPFC (**Fig. S40**). In Hippocampal GRN, SNP rs71350303 disrupts RELA regulation of TNFAIP6 (**Fig. S41B**), and in the LTL GRN, SNP rs6425995 disrupts NFKB1 regulation of CSF3R **(Fig. S41C**).

Besides the NFKB pathway, we also found several other shared pathways in AD and Covid-19 from KEGG, such as IKK, TNFR, PI3K, JNK, and IL6 (**Fig. S42**). We thus looked at the correlations between AD phenotypes and genes from those pathways in each brain region (**Fig. S43**). Finally, we identified highly correlated AD-COVID pathways and AD phenotypes (e.g., TNFR with severe stage and IKK with cognitive impairment in Hippocampus, IKK with frequent plaques in LTL, JNK with Resilience in DLPFC). Therefore, our integrative analysis helps identify Covid-19 susceptibility non-coding SNPs, which are also SNPs that worsen AD phenotypes, most likely via neuroinflammatory effects.

### Machine learning prediction of Covid-19 severity from AD-Covid gene regulatory networks

All gene lists and machine learning prediction results for this section are available in **Supp. File 9**. In total, 22 genes are shared between Covid-19 and AD KEGG database pathways. We also looked at our brain-region GRNs related to those AD-Covid genes, including TFs that regulate them and their target genes, i.e., AD-Covid GRNs. We found 1,305 genes from Hippocampus GRN, 670 genes from LTL GRN, and 895 genes from DLPFC GRN (2,536 unique genes in total, and 38 genes shared by three regions). As shown in **Fig. S44**, those 38 shared genes (17 found in both Covid and AD KEGG Pathways) across AD-Covid GRNs also highly correlate with AD phenotypes such as clinical Braak stage.

764 out of 2,536 unique genes across our AD-Covid GRNs are significantly differentially expressed in the Covid-19 severity condition. Moreover, the Covid-19 phenotype is positively correlated with many mechanisms in the AD KEGG pathway (**Fig. S45**). We normalized gene expression data (**Fig. S46**) of a recent Covid-19 cohort (N=50 ICU vs. N=50 non-ICU)^108^ and identified differentially expressed genes (DEGs) for Covid-19 ICU (**Methods**). We found 5,085 DEGs (2,505 up-regulated, 2,580 down-regulated genes) in total. Of those DEGs, 460 DEGs are from the AD-Covid GRN in Hippocampus (240 up-regulated, 220 down-regulated, **Fig. 6A**). Similarly, LTL’s AD-Covid GRN has 164 DEGs (88 up-regulated, 76 down-regulated, **Fig. 6B**). DLPFC’s has 254 DEGs (126 up-regulated, 128 down-regulated, **Fig. 6C**). 18 differentially expressed genes (13 up-regulated, 5 down-regulated), including SPI1 (up-regulated in Covid ICU patients), were found in AD-Covid GRNs for all 3 brain regions. These DEGs suggest that genes from our AD-Covid GRNs significantly associate with Covid-19 severity. Thus, beyond association, we further want to develop a model to predict Covid-19 severity from those genes. We did not use age and gender as predictors as both variables had a relatively low correlation with Covid-19 severity (r = 0.0943 for age, r = 0.0824 for gender).

**Fig 6.**
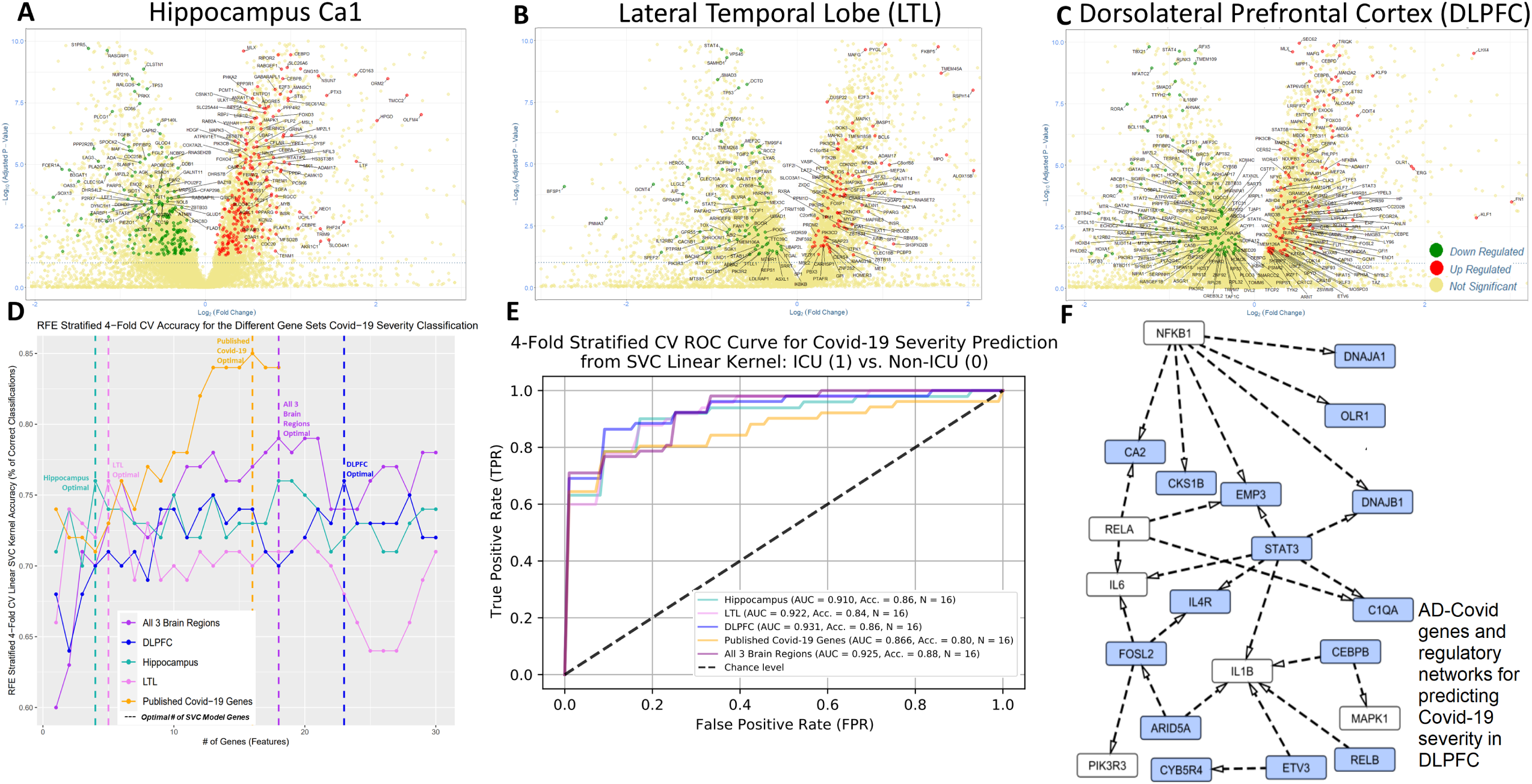
Differential expression and prediction of Covid-19 severity using AD-Covid gene regulatory networks. Top volcano plots show top differential expression analyses for Covid-19 severity (i.e., Covid-19 ICU, **Methods**) and label the genes from the AD-Covid gene regulatory networks (GRNs) that relate to 22 AD-Covid genes: (**A**) Hippocampus, (**B**) Lateral Temporal Lobe (LTL), (**C**) Dorsolateral Prefrontal Cortex. Red: up-regulated. Green: down-regulated. Yellow: No significance. x-axis: log2(fold change). y-axis: -log10(Adjusted p-value). (**D**) Prediction accuracy of Covid-19 severity after selecting different numbers of genes from AD-Covid GRNs and recently found Covid-19 genes (benchmark genes). The accuracy was calculated based on the support vector machine classification with 4-fold cross-validation. The dashed lines correspond to the minimal numbers of select genes with highest accuracy (i.e., optimal gene sets for predicting Covid-19 severity). (**E**) Receiver Operating Characteristic (ROC) curves and area under curve (AUC) values for classifying Covid-19 severity in (**D**). (**F**) Subnetwork of DLPFC GRN relating to the 16 DLPFC optimal genes (excluding MAPK14) for predicting Covid-19 severity (N=16) with AD-Covid shared genes. Blue: genes/TFs found in the DLPFC final model. White: AD-Covid shared genes. There was no overlap between both sets.

In particular, we first gathered 18 benchmark Covid-19 susceptibility genes from recent studies^50–53^. Then, we used recursive feature elimination (RFE) to select an optimal number of genes from the list of 18 published Covid-19 genes for Covid-19 severity, i.e., 4-fold stratified cross-validated feature selection with highest classification accuracy based on linear support vector machine (**Methods, Figures 6D** and **S47**). 16 genes were optimal for the published genes model. Then, we performed RFE feature selection for each region’s AD-Covid GRN to select the top 16 genes; building linear kernel SVM predicted probability models with the same number of genes (16) from each of the 5 gene lists would enable us to directly compare the effectiveness of our AD-Covid GRNs with that of the published genes (benchmark).

Our prediction accuracy based on 16 genes for each AD-Covid GRN is higher than the optimal benchmark of 16 Covid-19 genes (86% for Hippocampus, 86% for DLPFC, 84% for LTL, and 88% for combined regions, 80% for benchmark). As shown in **Fig. 6E**, our areas under the ROC curve (AUROC) values are larger than the benchmark (0.910 for Hippocampus, 0.931 for DLPFC, 0.922 for LTL, 0.925 for combined regions, 0.866 for benchmark). We show ROC curves for each of the 4 folds for all 5 models in **Fig. S48**. Relative to the benchmark, the optimal model (in terms of the highest accuracy and highest AUC, the model of the combined region) improved accuracy by 8% and boosted the AUC by 0.059. Therefore, the select genes from our AD-Covid GRNs have higher predictability than existing Covid-19 genes for predicting Covid-19 severity (and, by extension, predicting immune system dysregulation).

AD-Covid Genes driving Covid-19 severity can also drive neuroinflammation. 66 unique genes were selected across all 5 different models (51 were in at least 1 of our 4 AD-Covid GRNs). ADAM17 was the only AD-Covid gene found in the final gene list (Hippocampus model). 4 genes were found in 3 gene lists: CCR2 for combined, Hippocampus, benchmark models; CD58 and PLAUR for combined, Hippocampus, LTL models; and CEBPB for combined, DLPFC, Hippocampus models. CEBPB is a key TF for brain inflammation that binds to regulatory regions found on acute-phase and cytokine genes^97^. CEBPB regulates IL1B and MAPK1 in the DLPFC (**Fig. 6E**) and OLFM4, HMOX1, and TNFRSF1A (belongs to Severe Stage greenyellow Hippocampal gene module, associated with various abnormal immunological enrichments (**Fig. S8**)) in Hippocampus (**Fig. S49**). These 6 genes were found in 2 gene lists: CYB5R4 (protects cells from excess ROS) for DLPFC and LTL; PDK4 (metabolic regulator of resistance iron-dependent programmed cell death) for LTL and Hippocampus; DNAJB1, MAPK14, and CA2 for combined and DLPFC; OLFM4 (previously ranked top 26 proteins for AD based on a study using multi-omic analysis for proteomes^99^) for combined and Hippocampus models.

We highlight the subnetwork of the DLPFC GRN for 15 out of the 16 optimal predictive genes that are directly regulating or regulated by at least 1 of the 22 shared AD-Covid genes (**Fig. 6F**); we do the same for the Hippocampal GRN for 7 out of the 16 optimal predictive genes, excluding MAPK14 in **Fig. S49** and the LTL GRN for NFKB1 regulating PLAUR in **Fig. S50**. These brain region-specific subnetworks reveal TF-TG interactions that may predict levels of pro-inflammatory cytokines and neuroinflammation (severity of immune system dysregulation based on Covid-19 severity levels). NFKB1 regulates 6 genes and RELA regulates 2 genes found in the final DLPFC model, further underscoring the importance of NF-kB TFs in regulating genes associated with immune dysregulation in DLPFC (based on abilities to predict Covid-19 severity levels). CA2 and EMP3 (up-regulated in glioma tissues^100^) are jointly regulated by RELA and NFKB1 TFs. RELB regulates IL1B in DLPFC. OLR1 has been previously identified as a candidate AD gene, and our results verify its potential role in regulating inflammation in AD.

CREB1, which targets PIK3R3 and PDK4 in our Hippocampus sub-network (**Fig. S49**), is implicated as a key transcriptional regulator in synaptic plasticity, learning and memory, and found to cross-talk with NFKB TFs^96^. RELA regulates NFKB1, which regulates 2 Hippocampal model genes. Moreover, 3 genes (PIK3R1, PIK3CA, PIK3R3) are regulatory subunits of PI3K, a shared AD-Covid mechanism; altered signaling of the PI3K-Akt insulin pathway in Type 2 Diabetes is associated with not only AD progression^101^, but also with worse Covid-19 outcomes via increased IRF5 activity in both diseases^59^. All in all, our AD-Covid genes may be biologically meaningful markers for a dysregulated immune system response in AD.

We also investigated the role of AD SNPs in regulating these AD-Covid genes, these potential AD biomarker genes for neuroinflammation. In **Fig. S50**, we found SNPs that may disrupt TF regulation of target genes within our Hippocampal network. For example, rs192376432 strongly disrupts CEBPB regulation of TNFRSF1A, which belongs to severe stage AD greenyellow module in the Hippocampus with many meaningful biological enrichments (**Fig. S8)**. SNPs rs761726261 and rs148956807 strongly disrupt CEBPB regulation of HMOX2 (correlated with Braak 3 (r = 0.383) and belongs to a Control Hippocampal module). At the same time, HMOX2 has been considered a potential risk gene in AD pathogenesis (with decreased levels found in AD patients and mice), previous studies^102^, were unable to detect any association between the coding SNPs in HMOX2 and AD. Our pipeline has uncovered several non-coding SNPs in the Hippocampus, with p < 8e-05, such as rs537044906 (disrupts RFX2 regulation of HMOX2) and rs559787069 (disrupts SPI1 regulation of HMOX2). Furthermore, rs559787069 is located between peak signals across all cell types, including microglia (**Fig. S51**). This is another example of how our integrative analysis can provide more insights on AD risk genes and SNPs that may potentially drive their expression changes in AD.

We also evaluated and compared our predictive models with benchmark genes using Decision Curve Analysis (DCA). DCA enables evaluating the clinical usability of our Covid-19 severity prediction models based on their Net Benefits (**Methods**). We plotted the Decision Curves to show how the Net Benefit of each model varies across probability thresholds (**Fig. S52**). And the threshold of each model that gives the highest Net Benefit corresponds to the optimal decision probability for sending Covid-19 patients to ICU or not, i.e., the “optimal” threshold (**Fig. S53**). In general, the models of genes from our AD-Covid GRNs have higher Net Benefits than the benchmark Covid-19 genes, especially around the optimal thresholds that achieve possible maximum Net Benefits. Note that since 50% of our Covid-19 positive patients are in the ICU, the maximum Net Benefit is 0.50. The increase in Net Benefit of our models (around 0.0011 to 0.247 at optimal thresholds with an average increase of 0.118 in Net Benefit) compared with the benchmark could be interpreted that using the genes from our AD-Covid GRNs on average increases the number of truly severe Covid-19 patients detected by about 118 per 1,000 Covid-19 positive patients, without changing the number of non-severe patients who are needlessly sent to the ICU (**Fig. S54**). The benchmark model was not optimal for any of the probability thresholds. Thus, those genes, along with our predictive models, provide potential novel strategies for helping clinical decisions on sending Covid-19 patients to ICU or not and may serve as candidates for neuroimmunology in AD.

## Discussion

This paper used 3 AD-relevant brain regions to uncover molecular mechanisms associated with AD progression in 3 AD-relevant brain regions (DLPFC, Hippocampus, LTL). We linked disease variants to TFs to regulatory elements to genes and modules. Further, we investigated neuroimmunology pathways targeted by AD by utilizing Covid-19 severity (a proxy for dysregulated immunity), determining potential biomarkers for neuroinflammation. AD progression is linked to immunological mechanisms in the brain; pattern recognition receptors on microglia and astrocytes may respond to misfolded aggregated proteins in the brain, releasing inflammatory cytokines, driving AD progression^103^. Studies emphasize systems biology approaches (like ours) to identify biomarkers to target AD-associated neuroinflammation^88^; AD itself has over 34 canonical intricately interconnected pathways, making that process daunting^104^. Focusing on shared AD-Covid pathways may be a helpful point of departure; AD and Covid-19 have strong links and exacerbate one another. We found non-coding SNPs impacting neuroimmunology pathways correlated with AD phenotypes; these SNPs may help explain genetic mechanisms of critical illness in Covid-19^51^, and be possible candidate loci for neuroimmunology in AD.

Our findings help specifically support further research into the links between shared AD and Covid-19 pathways and the role of pro-inflammatory cytokines and rogue immune system responses in AD^8^. Perhaps common AD-Covid pathways could be targeted through treatments to alleviate the suffering of patients with either disease; for instance, researchers could explore suppressing Interferon response or stimulating the cholinergic anti-inflammatory pathway in Covid-19 patients acetylcholinesterase inhibitors (a current AD treatment strategy)^59^.

Nonetheless, more research is needed to verify the true causal links between Covid-19 and AD. There are also some limitations to our work. Brain regions are composed of different cell types that may impact co-expression networks and gene regulation; for example, AD patients may have fewer neurons and more immune cells. Recent single-cell sequencing data (e.g., scRNA-seq, scATAC-seq) in AD enable studying functional genomics and regulatory mechanisms at cell type levels^105^. Many cell-type GRNs in the human brain, such as neurons and glia, were predicted from single-cell data. Our paper validated many phenotype-associated SNPs locate on regulatory elements with cell-type epigenomic activities, suggesting potential cell-type regulatory effects of those SNPs. We can perform an integrative analysis of cell-type networks to understand regulatory mechanisms from variants that cause AD for various cell types. GWAS studies identified additional variants associated with refined AD phenotypes such as cerebrospinal fluid and psychotic symptoms and many genetic variants associated with neuroinflammation^88^. We aim to predict GRNs of those variants for additional AD phenotypes in the future.

Covid-19 patient transcriptomic data was collected from blood samples and may present limitations, especially since AD-Covid GRNs are based on transcriptomic data from brain region samples. Nonetheless, researchers have found signs of immune dysregulation in both AD brain and blood samples^106^. Elevated chemokines and proinflammatory molecules in Covid-19 patients can compromise the Blood Brain Barrier (breaks down in AD) and enter the brain, encountering astrocytes and microglia (malfunctioning in AD); such patients are more susceptible to severe Covid-19 outcomes and further neurological damage. Thus, Covid-19 patient blood samples data may potentially predict future CNS invasion and neuroinflammation; we thus used this blood-brain link to motivate our machine learning application, where we tried to uncover biomarkers associated with the immune system dysregulation during AD.

Many large scientific consortia have generated matched multi-omic data of individuals such as PsychENCODE^31^, AMP-AD^107^, TCGA^108^. Moreover, machine learning has been widely used for predicting phenotypes from individual data. Thus, we can extend our analysis to input matched omics data of individuals (e.g., genotype, gene expression, epigenomics) at the population level to train machine learning models to predict personalized phenotypes. The resulting predictive models can further predict personalized phenotypes for new individual data and prioritize phenotype-specific functional genomics and GRNs in human diseases. In addition, we found TFs regulate multiple gene co-expression modules and further link to different phenotypes. This suggests possible collinearity driven by those TF regulations across phenotypes. Emerging advanced machine learning approaches such as neural networks will potentially decouple such phenotypic collinearity and help find phenotypic-specific TFs.

Neuroimmunology research is uncovering the role of dysregulated and sustained immune responses in other complex neurologic diseases, such as Schizophrenia, Myasthenia Gravis, Parkinson’s disease, Amyotrophic Lateral Sclerosis, and Multiple Sclerosis^109^. Overall, we hope that our approach to analyzing molecular mechanisms in AD can be applied to help understand molecular mechanisms associated with other diseases by uncovering the association of orphaned GWAS loci in non-coding regions with disease phenotypes and using other closely related diseases to help reveal additional mechanisms at play. For instance, Covid-19 can be a proxy to understand the role of a dysregulated immune system in Schizophrenia, especially since Schizophrenia is an autoimmune disease (excessive synaptic pruning by microglia) and is the second-largest risk factor for Covid-19 death^110^.

## Conclusion

We performed integrative multi-omics analysis from genotype, chromatin interactions, transcriptomics, and epigenomics to predict gene regulatory networks (GRNs) for various AD phenotypes in three major AD-related brain regions: Hippocampus, Dorsolateral Prefrontal Cortex, and Lateral Temporal Lobe. Those brain-region GRNs link AD risk variants (i.e., SNPs) to disrupted binding sites of transcription factors to regulatory elements (e.g., enhancers, promoters) to target genes or gene co-expression modules. Our comparative network analyses further revealed cross-brain and brain-specific gene regulatory network structures from risk SNPs to phenotypes in AD. Many phenotype-associated SNPs locate on regulatory elements that have cell-type epigenomic activities, suggesting potential cell-type regulatory effects of those SNPs. We used potential gene regulatory connections between AD and Covid-19 to further investigate the role of neuroimmunology in AD. Our machine learning analysis further prioritized genes for predicting severe Covid as a set of “AD-Covid genes,” providing potentially novel gene biomarkers between Covid-19, neuroimmunology, and AD. These genes that predict Covid-19 severity are thus possible biomarkers for indicating immune system dysregulation and inflammation levels in AD. Decision Curve Analysis (DCA) showed that our AD-Covid genes outperform known Covid-19 genes for predicting Covid severity and making the decision to send Covid patients to ICU, demonstrating clinical translational ability of our predictive models. Using our GRNs, we reported AD SNPs linked to those AD-Covid genes, providing possible genetic causes for dysregulation of neuroimmunology in AD-Covid. Our analysis is open-source available and thus can serve as general purpose for understanding functional genomics and gene regulation for other diseases. Finally, all our analysis (codes and data) is open source at https://github.com/daifengwanglab/ADSNPheno for reproducibility and general usage, and a web database at https://adsnpheno.shinyapps.io/AlzheimersDisease_SNPheno for visualizing our findings.

## Supplementary Information

Supplementary file 1 – Genes, modules, phenotypes, enrichments, and Transcription Factors regulating gene modules for Hippocampus CA1

Supplementary file 2 – Genes, modules, phenotypes, and enrichments and Transcription Factors regulating gene modules for Lateral Temporal Lobe (LTL)

Supplementary file 3 – Genes, modules, phenotypes, and enrichments for the Dorsolateral Prefrontal Cortex (DLPFC)

Supplementary file 4 – Filtered Gene Regulatory Network (GRN) for the Hippocampus CA1 (based on target genes associated with AD phenotypes or belonging to modules associated with AD phenotypes)

Supplementary file 5 – Gene Regulatory Network for the Lateral Temporal Lobe (LTL)

Supplementary file 6 – SNPs Interrupting Transcription Factor Binding Sites (TFBS) in Hippocampus Ca1 Gene Regulatory Network (GRN)

Supplementary file 7 – SNPs Interrupting Transcription Factor Binding Sites (TFBS) in Lateral Temporal Lobe (LTL) Gene Regulatory Network (GRN)

Supplementary file 8 – SNPs Interrupting Transcription Factor Binding Sites (TFBS) in DLPFC Gene Regulatory Network (GRN)

Supplementary file 9 – shared AD and Covid-19 genes, 5 initial gene lists (for Hippocampus Ca1, Lateral Temporal Lobe (LTL), Dorsolateral Prefrontal Cortex (DLPFC), All 3 combined, Published Covid-19 genes), Differential Expression Analysis genes found in AD-COVID GRNs for each region, final SVC linear kernel genes for 5 models, predicted probabilities from each model, and Decision Curve Analysis (Net Benefit of each model for various probability thresholds).

Supplementary document – Supplementary Figures S1 to S54 and Supplementary Tables S1 to S3.

## Acknowledgments

The authors are grateful to Jonathan Edward Bryan for his assistance in tasks related to this project and Renu Poochie Khullar for her helpful comments on the manuscript.

## Funding

This work was supported by the grants of National Institutes of Health, R01AG067025, R21CA237955 and U01MH116492 to Daifeng Wang and U54HD090256 to Waisman Center. Saniya Khullar was supported by an NLM training grant to the Computation and Informatics in Biology and Medicine Training Program (NLM 5T15LM007359).

## Competing Interests

None declared.

## Authors’ Contributions

D.W. conceived and designed the study. S.K. and D.W. analyzed the data and wrote the manuscript.

## Ethics approval and consent to participate

Not applicable. Public data was utilized for this analysis as well as approved ROSMAP data for the DLPFC.

## Consent for publication

All authors read and approved the final manuscript.

## Availability of data and material

Our analysis is open-source available at https://github.com/daifengwanglab/ADSNPheno and our functional genomics resource for AD is available at https://adsnpheno.shinyapps.io/AlzheimersDisease_SNPheno.

## Notes

### Competing Interest Statement

The authors have declared no competing interest.

## References

1. Alzheimer’s Statistics. Alzheimers.net https://www.alzheimers.net/resources/alzheimers-statistics.

2. Rabinovici, G. D. Late-onset Alzheimer Disease. Continuum (Minneap Minn) 25, 14–33 (2019).

3. Morabito, S., Miyoshi, E., Michael, N. & Swarup, V. Integrative genomics approach identifies conserved transcriptomic networks in Alzheimer’s disease. Human Molecular Genetics 29, 2899–2919 (2020).

4. Co-expression modules construction by WGCNA and identify potential prognostic markers of uveal melanoma - ScienceDirect. https://www.sciencedirect.com/science/article/abs/pii/S0014483517303536?via%3Dihub.

5. Jansen, I. E. et al. Genome-wide meta-analysis identifies new loci and functional pathways influencing Alzheimer’s disease risk. Nat Genet 51, 404–413 (2019).

6. Novikova, G. et al. Integration of Alzheimer’s disease genetics and myeloid genomics identifies disease risk regulatory elements and genes. Nature Communications 12, 1610 (2021).

7. Wingo, A. P. et al. Integrating human brain proteomes with genome-wide association data implicates new proteins in Alzheimer’s disease pathogenesis. Nature Genetics 53, 143–146 (2021).

8. Kinney, J. W. et al. Inflammation as a central mechanism in Alzheimer’s disease. Alzheimers Dement (N Y) 4, 575–590 (2018).

9. Comprehensive functional genomic resource and integrative model for the human brain | Science. https://science.sciencemag.org/content/362/6420/eaat8464.

10. Nuclear factor-kappa β as a therapeutic target for Alzheimer’s disease - Jha - 2019 - Journal of Neurochemistry - Wiley Online Library. https://onlinelibrary.wiley.com/doi/10.1111/jnc.14687.

11. Amruta, N. et al. SARS-CoV-2 mediated neuroinflammation and the impact of COVID-19 in neurological disorders. Cytokine & Growth Factor Reviews 58, 1–15 (2021).

12. Neurological Manifestations of COVID-19 Feature T Cell Exhaustion and Dedifferentiated Monocytes in Cerebrospinal Fluid: Immunity. https://www.cell.com/immunity/fulltext/S1074-7613(20)30539-2?_returnURL= https://linkinghub.elsevier.com%2Fretrieve%2Fpii%2FS1074761320305392%3Fshowall%3Dtrue.

13. Erausquin, G. A. de et al. The chronic neuropsychiatric sequelae of COVID-19: The need for a prospective study of viral impact on brain functioning. Alzheimer’s & Dementia 17, 1056–1065 (2021).

14. COVID-19 Tied to Acceleration of Alzheimer’s Pathology. Medscape http://www.medscape.com/viewarticle/955755.

15. Inal, J. Biological Factors Linking ApoE ε4 Variant and Severe COVID-19. Curr Atheroscler Rep 22, (2020).

16. steve. What is NF-κB pathway? MBInfo https://www.mechanobio.info/what-is-mechanosignaling/signaling-pathways/what-is-the-nf-%ce%bab-pathway/.

17. Su, C.-M., Wang, L. & Yoo, D. Activation of NF-κB and induction of proinflammatory cytokine expressions mediated by ORF7a protein of SARS-CoV-2. Sci Rep 11, 13464 (2021).

18. Janssen, N. A. F. et al. Dysregulated Innate and Adaptive Immune Responses Discriminate Disease Severity in COVID-19. J Infect Dis 223, 1322–1333 (2021).

19. Blalock, E. M. et al. Incipient Alzheimer’s disease: Microarray correlation analyses reveal major transcriptional and tumor suppressor responses. PNAS 101, 2173–2178 (2004).

20. http://mail.nih.gov>, S. D. <sdavis2 at. GEOquery: Get data from NCBI Gene Expression Omnibus (GEO). (Bioconductor version: Release (3.12), 2021). doi:10.18129/B9.bioc.GEOquery.

21. hgu133a.db. Bioconductor http://bioconductor.org/packages/hgu133a.db/.

22. hgu133acdf. Bioconductor http://bioconductor.org/packages/hgu133acdf/.

23. http://ds.harvard.edu>, R. A. I. <rafa at et al. affy: Methods for Affymetrix Oligonucleotide Arrays. (Bioconductor version: Release (3.12), 2021). doi:10.18129/B9.bioc.affy.

24. RMA normalization for microarray data. https://felixfan.github.io/RMA-Normalization-Microarray/.

25. Nativio, R. et al. An integrated multi-omics approach identifies epigenetic alterations associated with Alzheimer’s disease. Nature Genetics 52, 1024–1035 (2020).

26. The Religious Orders Study and Memory and Aging Project (ROSMAP) Study. https://adknowledgeportal.synapse.org/Explore/Studies/DetailsPage?Study=syn3219045.

27. An atlas of chromatin accessibility in the adult human brain. https://bendlj01.u.hpc.mssm.edu/multireg/.

28. Jung, I. et al. A compendium of promoter-centered long-range chromatin interactions in the human genome. Nat Genet 51, 1442–1449 (2019).

29. TxDb.Hsapiens.UCSC.hg19.knownGene. Bioconductor http://bioconductor.org/packages/TxDb.Hsapiens.UCSC.hg19.knownGene/.

30. Rödelsperger, C. et al. Short ultraconserved promoter regions delineate a class of preferentially expressed alternatively spliced transcripts. Genomics 94, 308–316 (2009).

31. Resource.PsychEncode. http://resource.psychencode.org/.

32. Langfelder, P. & Horvath, S. WGCNA: an R package for weighted correlation network analysis. BMC Bioinformatics 9, 559 (2008).

33. Botía, J. A. CoExpNets brief instructions. (2021).

34. Botía, J. A. et al. An additional k-means clustering step improves the biological features of WGCNA gene co-expression networks. BMC Systems Biology 11, 47 (2017).

35. Safieh, M., Korczyn, A. D. & Michaelson, D. M. ApoE4: an emerging therapeutic target for Alzheimer’s disease. BMC Medicine 17, 64 (2019).

36. Jin, T. et al. scGRNom: a computational pipeline of integrative multi-omics analyses for predicting cell-type disease genes and regulatory networks. Genome Medicine 13, 95 (2021).

37. Tan, G. TFBSTools: Software Package for Transcription Factor Binding Site (TFBS) Analysis. (Bioconductor version: Release (3.12), 2021). doi:10.18129/B9.bioc.TFBSTools.

38. Schep, A. & University, S. motifmatchr: Fast Motif Matching in R. (Bioconductor version: Release (3.12), 2021). doi:10.18129/B9.bioc.motifmatchr.

39. Groeneveld, C. et al. RTN: RTN: Reconstruction of Transcriptional regulatory Networks and analysis of regulons. (Bioconductor version: Release (3.12), 2021). doi:10.18129/B9.bioc.RTN.

40. http://systemsbiology.org>, S. A. <seth ament at, http://systemsbioloyg.org>, P. S. <pshannon at & http://systemsbiology.org>, M. R. <mrichard at. trena: Fit transcriptional regulatory networks using gene expression, priors, machine learning. (Bioconductor version: Release (3.12), 2021). doi:10.18129/B9.bioc.trena.

41. Huynh-Thu, V. A., Aibar, S. & Geurts, P. GENIE3: GEne Network Inference with Ensemble of trees. (Bioconductor version: Release (3.12), 2021). doi:10.18129/B9.bioc.GENIE3.

42. The Human Transcription Factors: Cell. https://www.cell.com/cell/fulltext/S0092-8674(18)30106-5?_returnURL= https://linkinghub.elsevier.com%2Fretrieve%2Fpii%2FS0092867418301065%3Fshowall%3Dtrue.

43. JASPAR 2020: An open-access database of transcription factor binding profiles. http://jaspar.genereg.net.

44. Margolin, A. A. et al. ARACNE: An Algorithm for the Reconstruction of Gene Regulatory Networks in a Mammalian Cellular Context. BMC Bioinformatics 7, S7 (2006).

45. Coetzee, S. G. & Hazelett, D. J. motifbreakR: A Package For Predicting The Disruptiveness Of Single Nucleotide Polymorphisms On Transcription Factor Binding Sites. (Bioconductor version: Release (3.12), 2021). doi:10.18129/B9.bioc.motifbreakR.

46. KEGG PATHWAY Database. https://www.genome.jp/kegg/pathway.html.

47. Luo, W. pathview: a tool set for pathway based data integration and visualization. (Bioconductor version: Release (3.12), 2021). doi:10.18129/B9.bioc.pathview.

48. Overmyer, K. A. et al. Large-Scale Multi-omic Analysis of COVID-19 Severity. Cell Syst 12, 23–40.e7 (2021).

49. Love, M., Ahlmann-Eltze, C., Forbes, K., Anders, S. & Huber, W. DESeq2: Differential gene expression analysis based on the negative binomial distribution. (Bioconductor version: Release (3.12), 2021). doi:10.18129/B9.bioc.DESeq2.

50. Hu, J., Li, C., Wang, S., Li, T. & Zhang, H. Genetic variants are identified to increase risk of COVID-19 related mortality from UK Biobank data. Human Genomics 15, 10 (2021).

51. Genetic mechanisms of critical illness in COVID-19 | Nature. https://www.nature.com/articles/s41586-020-03065-y#citeas.

52. New insights into genetic susceptibility of COVID-19: an ACE2 and TMPRSS2 polymorphism analysis | BMC Medicine | Full Text. https://bmcmedicine.biomedcentral.com/articles/10.1186/s12916-020-01673-z#citeas.

53. Kong, Y. et al. VEGF-D: a novel biomarker for detection of COVID-19 progression. Critical Care 24, 373 (2020).

54. scikit-learn: machine learning in Python — scikit-learn 0.24.1 documentation. https://scikit-learn.org/stable/.

55. Vickers, A. J. & Elkin, E. B. Decision Curve Analysis: A Novel Method for Evaluating Prediction Models. Med Decis Making 26, 565–574 (2006).

56. Biostatistics: Decision Curve Analysis | Memorial Sloan Kettering Cancer Center. https://www.mskcc.org/departments/epidemiology-biostatistics/biostatistics/decision-curve-analysis.

57. CA1 neurons in the human hippocampus are critical for autobiographical memory, mental time travel, and autonoetic consciousness | PNAS. https://www.pnas.org/content/108/42/17562.

58. Lee, J. K. & Kim, N.-J. Recent Advances in the Inhibition of p38 MAPK as a Potential Strategy for the Treatment of Alzheimer’s Disease. Molecules 22, (2017).

59. Naughton, S. X., Raval, U. & Pasinetti, G. M. Potential Novel Role of COVID-19 in Alzheimer’s Disease and Preventative Mitigation Strategies. J Alzheimers Dis 76, 21–25 (2020).

60. JCI - Type I interferon response drives neuroinflammation and synapse loss in Alzheimer disease. https://www.jci.org/articles/view/133737.

61. Metabolic correlates of prevalent mild cognitive impairment and Alzheimer’s disease in adults with Down syndrome - Mapstone - 2020 - Alzheimer’s & Dementia: Diagnosis, Assessment & Disease Monitoring - Wiley Online Library. https://alz-journals.onlinelibrary.wiley.com/doi/full/10.1002/dad2.12028.

62. Wnt signaling in the nervous system and in Alzheimer’s disease | Journal of Molecular Cell Biology | Oxford Academic. https://academic.oup.com/jmcb/article/6/1/64/874321.

63. Neurodegeneration and microtubule dynamics: death by a thousand cuts. https://www.ncbi.nlm.nih.gov/pmc/articles/PMC4563776/.

64. Vasile, F., Dossi, E. & Rouach, N. Human astrocytes: structure and functions in the healthy brain. Brain Struct Funct 222, 2017–2029 (2017).

65. Prion protein and Alzheimer disease. https://www.ncbi.nlm.nih.gov/pmc/articles/PMC2807690/.

66. Brinton, R. D., Gore, A. C., Schmidt, P. J. & Morrison, J. H. 68 - Reproductive Aging of Females: Neural Systems. in Hormones, Brain and Behavior (Second Edition) (eds. Pfaff, D. W., Arnold, A. P., Etgen, A. M., Fahrbach, S. E. & Rubin, R. T.) 2199–2224 (Academic Press, 2009). doi:10.1016/B978-008088783-8.00068-1.

67. Kumar, S. et al. Dorsolateral prefrontal cortex metabolites and their relationship with plasticity in Alzheimer’s disease. Alzheimer’s & Dementia 16, e045879 (2020).

68. Frontiers | Microglia in Alzheimer Disease: Well-Known Targets and New Opportunities | Frontiers in Aging Neuroscience. https://www.frontiersin.org/articles/10.3389/fnagi.2019.00233/full.

69. Ebanks, B., Ingram, T. L. & Chakrabarti, L. ATP synthase and Alzheimer’s disease: putting a spin on the mitochondrial hypothesis. Aging (Albany NY) 12, 16647–16662 (2020).

70. Cho, S.-J., Park, M. H., Han, C., Yoon, K. & Koh, Y. H. VEGFR2 alteration in Alzheimer’s disease. Scientific Reports 7, 17713 (2017).

71. Frontiers | Therapeutic Inhibition of the Complement System in Diseases of the Central Nervous System | Immunology. https://www.frontiersin.org/articles/10.3389/fimmu.2019.00362/full.

72. Quintela-López, T. et al. Aβ oligomers promote oligodendrocyte differentiation and maturation via integrin β1 and Fyn kinase signaling. Cell Death & Disease 10, 1–16 (2019).

73. Kumar, S., Ambrosini, G. & Bucher, P. SNP2TFBS – a database of regulatory SNPs affecting predicted transcription factor binding site affinity. Nucleic Acids Res 45, D139–D144 (2017).

74. Lu, T. et al. REST and stress resistance in ageing and Alzheimer’s disease. Nature 507, 448–454 (2014).

75. Maezawa, I., Jenkins, D. P., Jin, B. E. & Wulff, H. Microglial KCa3.1 Channels as a Potential Therapeutic Target for Alzheimer’s Disease. Int J Alzheimers Dis 2012, (2012).

76. Magno, L. et al. Alzheimer’s disease phospholipase C-gamma-2 (PLCG2) protective variant is a functional hypermorph. Alzheimer’s Research & Therapy 11, 16 (2019).

77. Liu, N. et al. Hippocampal transcriptome-wide association study and neurobiological pathway analysis for Alzheimer’s disease. PLOS Genetics 17, e1009363 (2021).

78. Yashiro, T. et al. A transcription factor PU.1 is critical for Ccl22 gene expression in dendritic cells and macrophages. Sci Rep 9, 1161 (2019).

79. PU.1 regulates Alzheimer’s disease-associated genes in primary human microglia | Molecular Neurodegeneration | Full Text. https://molecularneurodegeneration.biomedcentral.com/articles/10.1186/s13024-018-0277-1.

80. Early induction of c-Myc is associated with neuronal cell death - ScienceDirect. https://www.sciencedirect.com/science/article/abs/pii/S0304394011013899.

81. Sáez, M. E. et al. Genome Wide Meta-Analysis identifies common genetic signatures shared by heart function and Alzheimer’s disease. Sci Rep 9, 16665 (2019).

82. Yi, M. et al. The potassium channel KCa3.1 constitutes a pharmacological target for astrogliosis associated with ischemia stroke. J Neuroinflammation 14, (2017).

83. Kircheis, R. et al. NF-κB Pathway as a Potential Target for Treatment of Critical Stage COVID-19 Patients. Front. Immunol. 11, (2020).

84. Kwee, T. C. & Kwee, R. M. Chest CT in COVID-19: What the Radiologist Needs to Know. RadioGraphics 40, 1848–1865 (2020).

85. Targeting transcriptional regulation of SARS-CoV-2 entry factors ACE2 and TMPRSS2 | PNAS. https://www.pnas.org/content/118/1/e2021450118.

86. Tremblay, M.-E., Madore, C., Bordeleau, M., Tian, L. & Verkhratsky, A. Neuropathobiology of COVID-19: The Role for Glia. Frontiers in Cellular Neuroscience 14, 363 (2020).

87. Liu, M. et al. CXCL10/IP-10 in infectious diseases pathogenesis and potential therapeutic implications. Cytokine Growth Factor Rev 22, 121–130 (2011).

88. Hampel, H. et al. A Path Toward Precision Medicine for Neuroinflammatory Mechanisms in Alzheimer’s Disease. Frontiers in Immunology 11, 456 (2020).

89. Frontiers | Neutrophils in COVID-19 | Immunology. https://www.frontiersin.org/articles/10.3389/fimmu.2021.652470/full.

90. Landhuis, E. Could the immune system be key to Alzheimer’s disease? Knowable Magazine | Annual Reviews (2021) doi:10.1146/knowable-012921-3.

91. Fagone, P. et al. Transcriptional landscape of SARS-CoV-2 infection dismantles pathogenic pathways activated by the virus, proposes unique sex-specific differences and predicts tailored therapeutic strategies. Autoimmun Rev 19, 102571 (2020).

92. Parham, P. The Immune System. (Garland Science, 2014).

93. Hammad, A., Westacott, L. & Zaben, M. The role of the complement system in traumatic brain injury: a review. J Neuroinflammation 15, (2018).

94. Java, A. et al. The complement system in COVID-19: friend and foe? JCI Insight 5,.

95. Heffron, A. S. et al. The landscape of antibody binding in SARS-CoV-2 infection. BioRxiv 2020.10.10.334292 (2021) doi:10.1101/2020.10.10.334292.

96. Snow, W. M. & Albensi, B. C. Neuronal Gene Targets of NF-κB and Their Dysregulation in Alzheimer’s Disease. Frontiers in Molecular Neuroscience 9, 118 (2016).

97. Cardinaux, J. R., Allaman, I. & Magistretti, P. J. Pro-inflammatory cytokines induce the transcription factors C/EBPbeta and C/EBPdelta in astrocytes. Glia 29, 91–97 (2000).

98. Song, X. et al. PDK4 dictates metabolic resistance to ferroptosis by suppressing pyruvate oxidation and fatty acid synthesis. Cell Reports 34, 108767 (2021).

99. Bai, B. et al. Deep Multilayer Brain Proteomics Identifies Molecular Networks in Alzheimer’s Disease Progression. Neuron 105, 975–991.e7 (2020).

100. Gao, Y.-F. et al. PPIC, EMP3 and CHI3L1 Are Novel Prognostic Markers for High Grade Glioma. International Journal of Molecular Sciences 17, 1808 (2016).

101. Gabbouj, S. et al. Altered Insulin Signaling in Alzheimer’s Disease Brain – Special Emphasis on PI3K-Akt Pathway. Front Neurosci 13, 629 (2019).

102. Shibata, N., Ohnuma, T., Baba, H. & Arai, H. No Genetic Association between Polymorphisms of Heme Oxygenase 1 and 2 and Alzheimer’s Disease in a Japanese Population. DEM 27, 273–277 (2009).

103. Heneka, M. T. et al. Neuroinflammation in Alzheimer’s Disease. Lancet Neurol 14, 388–405 (2015).

104. Mizuno, S. et al. AlzPathway: a comprehensive map of signaling pathways of Alzheimer’s disease. BMC Syst Biol 6, 52 (2012).

105. Jiang, J., Wang, C., Qi, R., Fu, H. & Ma, Q. scREAD: A Single-Cell RNA-Seq Database for Alzheimer’s Disease. iScience 23, 101769 (2020).

106. Guo, Z. et al. Evaluation of Peripheral Immune Dysregulation in Alzheimer’s Disease and Vascular Dementia. Journal of Alzheimer’s Disease 71, 1175–1186 (2019).

107. AD Knowledge Portal -. https://adknowledgeportal.synapse.org/.

108. The Cancer Genome Atlas Program - National Cancer Institute. https://www.cancer.gov/about-nci/organization/ccg/research/structural-genomics/tcga (2018).

109. Coyle, P. K. Dissecting the Immune Component of Neurologic Disorders: A Grand Challenge for the 21st Century. Front Neurol 2, 37 (2011).

110. Schizophrenia Second Only to Age as Greatest Risk Factor for COVID-19 Death. NYU Langone News https://nyulangone.org/news/schizophrenia-second-only-age-greatest-risk-factor-covid-19-death.

